# Xylem perforation plate phenotypes affect water use and drought adaptation in maize (*Zea mays* L.)

**DOI:** 10.1101/2024.05.10.593543

**Authors:** Christopher F. Strock, Cody L. DePew, Jagdeep S. Sidhu, Tianyu Xu, Jonathan P. Lynch

**Author notes:** **Author for correspondence:** Jonathan P. Lynch.

## Abstract

- **Rationale**: Xylem morphology in annual monocots is important for water use strategies in many agronomically important species.
- **Methods:** We assess how xylem perforation plates affect water use strategies in maize (*Zea mays* L.) through *in silico* modeling, empirical studies under water deficit in controlled environments, and in the field.
- **Key Result**: Significant genotypic variation for the prominence and frequency of perforation plates was observed in maize germplasm. Perforation plate phenotypes had high heritability, were associated with several QTL, and were pleiotropic across leaves, aerial nodal roots, and subterranean nodal roots. Perforation plate phenotypes did not affect vulnerability to cavitation, but modeling predicted that they should affect axial water transport, which was supported by *in situ* measurements of root segments. Metaxylem vessel length was correlated with the rate of root elongation, root depth, and deep-water utilization in mesocosms. Under drought stress in the field, variation in xylem vessel length was associated with leaf roll, leaf temperature, transpiration, photosynthesis, and grain yield.
- **Main Conclusion:** Phenotypic variation for xylem perforation plate phenotypes in maize directly affects axial water conductance and is part of a pleiotropic syndrome with greater root elongation and deeper rooting that improves adaptation to water deficit stress.

## Introduction

Plants moderate the transport of water from roots to leaves through a variety of diverse mechanisms including stomatal gating (Hetherington & Woodward, 2003; Raven, 2014), ion-mediated flow regulation through pit membranes (Zwieniecki *et al*., 2001; Nardini *et al*., 2011), and morphological variation in anatomy (Tyree *et al*., 1994; Loepfe *et al*., 2007; Fan *et al*., 2009). Ultimately, the efficiency of water conduction, resistance to cavitation, and repair of embolisms depend primarily on the structural characteristics of xylem vessels. Consequently, xylem vessel morphology is strongly linked to ecological regimes, where overall plant design is closely cued to xylem function (Jansen *et al*., 2004; Carlquist, 2012).

Although the relationship between xylem morphology and water use strategies is obvious, the bulk of literature exploring this topic has been developed for perennial dicot species, and studies focusing on the xylem structure and function in annual monocots are less abundant (Carlquist, 2012; Aleman-Sancheschulz *et al*., 2020). While the principles of water transport through xylem apply to all vascular plants, the relationships between xylem morphology and water use strategies vary significantly among Tracheophytes. In contrast to dicots, monocots lack secondary growth and seasonal development of vasculature, while perennial species are subject to different constraints and opportunities with regard to water acquisition and use compared to annuals (Strock & Lynch, 2020). Research addressing the extent of intraspecific variation in xylem morphology in annual monocots is lacking, but essential for understanding the capacity and control of hydraulic conductance in many agronomically important species (Comas *et al*., 2013; Hacke *et al*., 2017).

Mature, empty xylem vessels provide the primary pathway for the transpiration stream and the pathway along these vessels through internodes and plexuses is shown to be uninterrupted by living cells (Shane *et al.,* 2000). Consequently, hydraulic resistance to axial flow through the xylem is largely considered to be a function of the inner diameter of the vessels, which can be approximated by the Hagen-Poiseuille equation. Nevertheless, this metric of hydraulic resistance calculated from vessel diameters does not consider the structure and abundance of perforation plates that likely affect the axial transport of water through xylem conduits, vulnerability to cavitation under water stress, and the dynamics of embolism repair.

In annual monocots like maize (*Zea mays*), mature metaxylem vessel elements are delineated by simple perforation plates (Fig. 1) (Carlquist, 2012). These plates form when metaxylem mature at the cessation of elongation, and lignified secondary cell wall is deposited on the lateral walls of the cell. At maturity, the central region of the shared primary wall and middle lamella between vessel elements is hydrolyzed, leaving a simple perforation plate composed of a lignified rim between vessel elements (Wang *et al*., 1994). These lignified rims of the simple perforation plates in mature xylem add a heterogenous topography to the otherwise smooth, lumen-facing walls (McCully *et al*., 2014).

**Figure 1.**
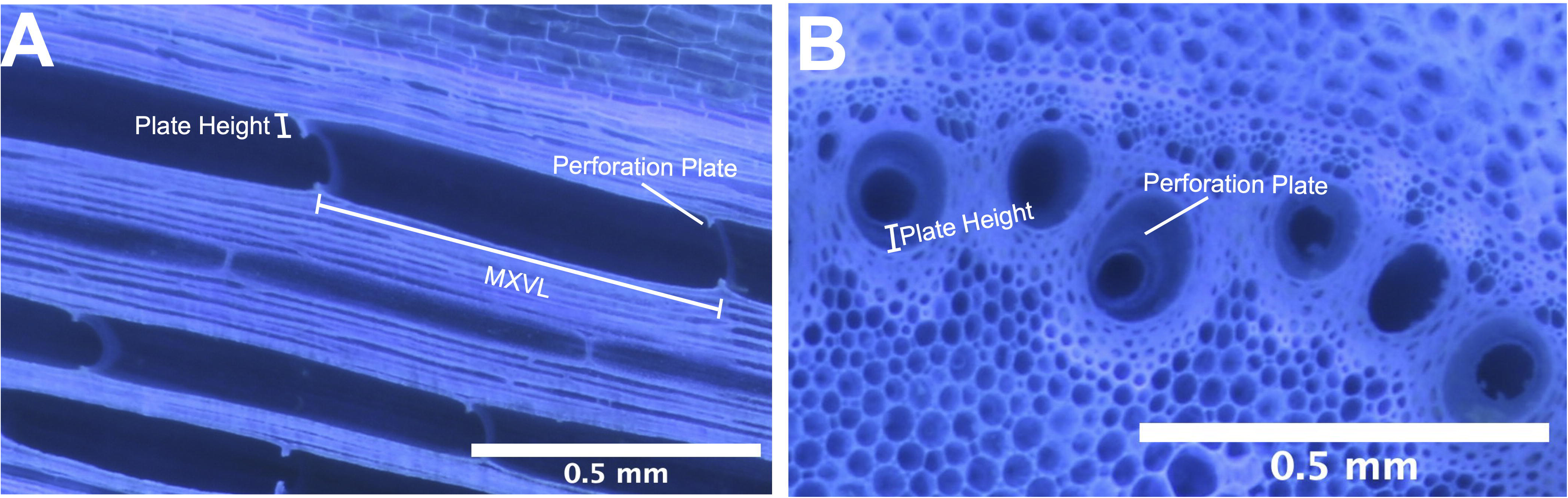
Longitudinal section and cross-section of maize (*Zea mays* L.) roots showing perforation plates within the metaxylem vessels.

Across taxa, the structure of these perforation plates is believed to be related to adaptive function in different environments, as differences in perforation plate characteristics are suggested to be optimally balancing trade-offs among conductive efficiency, mechanical strength, and resistance to cavitation. For example, the morphology of perforation plates has been shown to alter the relationship between the diameter of xylem vessels and plant height (Medeiros *et al*., 2019). Relationships between plate structure and climate have also been noted, where taxa with simple perforation plates that have less resistance to axial transport compared to other plate structures, tend to come from warm, dry climates where demand for hydraulic efficiency is essential (Jansen *et al*., 2004; Christman & Sperry, 2010).

While some studies suggest little to no added resistance to axial transport from simple perforation plates (Ellerby & Ennos, 1998; Zwieniecki *et al*. 2001; Sperry *et al*., 2006; Christman & Sperry, 2010), in other work, the Hagen-Poiseuille lumen resistivity is much less than the measured value, suggesting a significant effect of perforation plates on axial conductance (Sperry *et al*., 2005). For example, Gao *et al*., (2020) found that estimates of hydraulic conductance in xylem vessels of cotton calculated from the Hagen-Poiseuille equation were overestimated by more than 200%, largely due to the inability of the equation to represent perforation plates. Ultimately, the hydraulic resistance of these perforation plates is a function of their structure and the distance between successive plates along the length of the conduit, or the length of the vessel elements (Fig. 1) (Sperry *et al*., 2005).

In addition to their likely effect on axial transport, perforation plates act as pressure controllers that facilitate embolism repair (Lee *et al*., 2019). For example, Hwang *et al*., (2016) reports that in maize leaves, refilling of metaxylem vessels is halted at each perforation plate where refilling water then bypasses through protoxylem. This observation has implications for the hydrodynamic role of perforation plates to supply water into embolized vessels. Holbrook and Zwieniecki (1999) also suggest that perforation plates serve to isolate the embolism. *In vivo* visualization of embolized metaxylem in grape vine (*Vitis vinifera*) using microCT has revealed that water accumulates at the rim of simple perforation plates during refilling and perforation plates can act as loci for trapped gas bubbles in a refilling vessel (Brodersen *et al*., 2018). The stability of gas bubbles at the perforation plates of embolized vessels may suggest the lignified borders of perforation plates in *Vitis vinifera* are sufficiently hydrophobic (Brodersen *et al*., 2018). This variability in hydrophobicity of metaxylem cell walls has also been shown maize roots (McCully *et al*., 2014), and this makes sense as the hydrostatic connection to functional vessels would come with significant risk of water in the refilling vessel being pulled into the transpiration stream, leading to filling failures (Brodersen *et al*., 2010, 2018).

In this work, our goals were to explore the extent of intraspecific variation for the structure of perforation plates within an annual monocot and assess how this variation affects transport and use of water under drought stress. Specifically, we test the hypotheses that (1) simple perforation plates have a significant effect on water transport, (2) intraspecific variation exists for these features in maize, and (3) this variation affects water use strategies under drought stress. To test these hypotheses, we utilize *in silico* modeling as well as empirical observations of maize grown in controlled environment mesocosms and multiple field studies.

## Materials and Method

### Anatomical Measurements

To visualize internal anatomy of roots and leaves, samples were preserved, sectioned, and imaged by laser ablation tomography (LAT) as described in Strock *et al*., (2019, 2022). Analyzed dimensions of brace roots and subterranean roots include cross-sectional and longitudinal images. Analyzed dimensions of leaves include cross-sectional and longitudinal images of the lamina and the midrib, midway along the length of the leaf, approximately 2 cm from the midrib.

### Cryo-Scanning Electron Microscopy (SEM)

Cryo-SEM was used for closer observation of perforation plates. This work was completed using the Zeiss Sigma VP-FESEM at the Pennsylvania State University Huck Institutes of the Life Sciences Microscopy Core Facility. Samples of fourth node axial roots 10-14 cm from the base of the plant were collected at 37 d after planting and preserved in 75% (v/v) ethanol in water. The sample was mounted on a sample holder and plunged into liquid nitrogen. The sample holder was withdrawn under a vacuum into the cryo-preparation chamber where the sample was maintained at -195°C. The sample was then transferred to the SEM chamber onto a cold stage module. Variable pressure without sputter coating was used. Voltage was 10kV and samples were imaged at a temperature of -195°C.

### Germplasm Selection

For much of this work, we utilized maize recombinant inbred lines (RILs) from the intermated B73 x Mo17 (IBM) population. RILs descend from the same two parents, hence represent distinct genotypes sharing the same genetic background, thereby reducing the risk of confounding effects from genetic interactions, epistasis, and pleiotropy (Zhu *et al*., 2005, 2006). RILs are especially useful tools in cases in which the genetic basis of a phenotype is complex or unknown, as is the case with perforation plate prominence and metaxylem vessel length in maize, thereby precluding the use of single-gene variants.

To understand the variation that exists for perforation plate prominence and metaxylem vessel length, 234 RILs from the IBM population were grown at the Ukulima Root Biology Center (URBC) in Limpopo Province, South Africa (24.533367°S, 28.123783°E) from February through May 2011. At flowering, root crowns of two representative plants per genotype were extracted and washed as in Trachsel *et al*., (2011), and the anatomy was analyzed from one segment of nodal root from the fourth node of each plant, 5 cm from the base of the root. Root segments were sectioned and imaged using LAT as described above. The lengths of fifteen representative metaxylem vessels measured in LAT images of each root segment using imageJ (Schneider *et al*., 2012). From these 234 RILs of the IBM population, four accessions with distinct metaxylem vessel lengths were selected for use in subsequent controlled environment and field studies (Fig. 2). The four IBM accessions included IBM015 and IBM111, which were classified as having roots with less prominent (short) perforation plates and long metaxylem vessels, and IBM177 and IBM205, which were classified as having roots with prominent (tall) perforation plates and short metaxylem vessels. Seeds used in all glasshouse and field studies were provided by Dr. Shawn M. Kaeppler from U-Wisconsin, Madison, USA.

**Figure 2.**
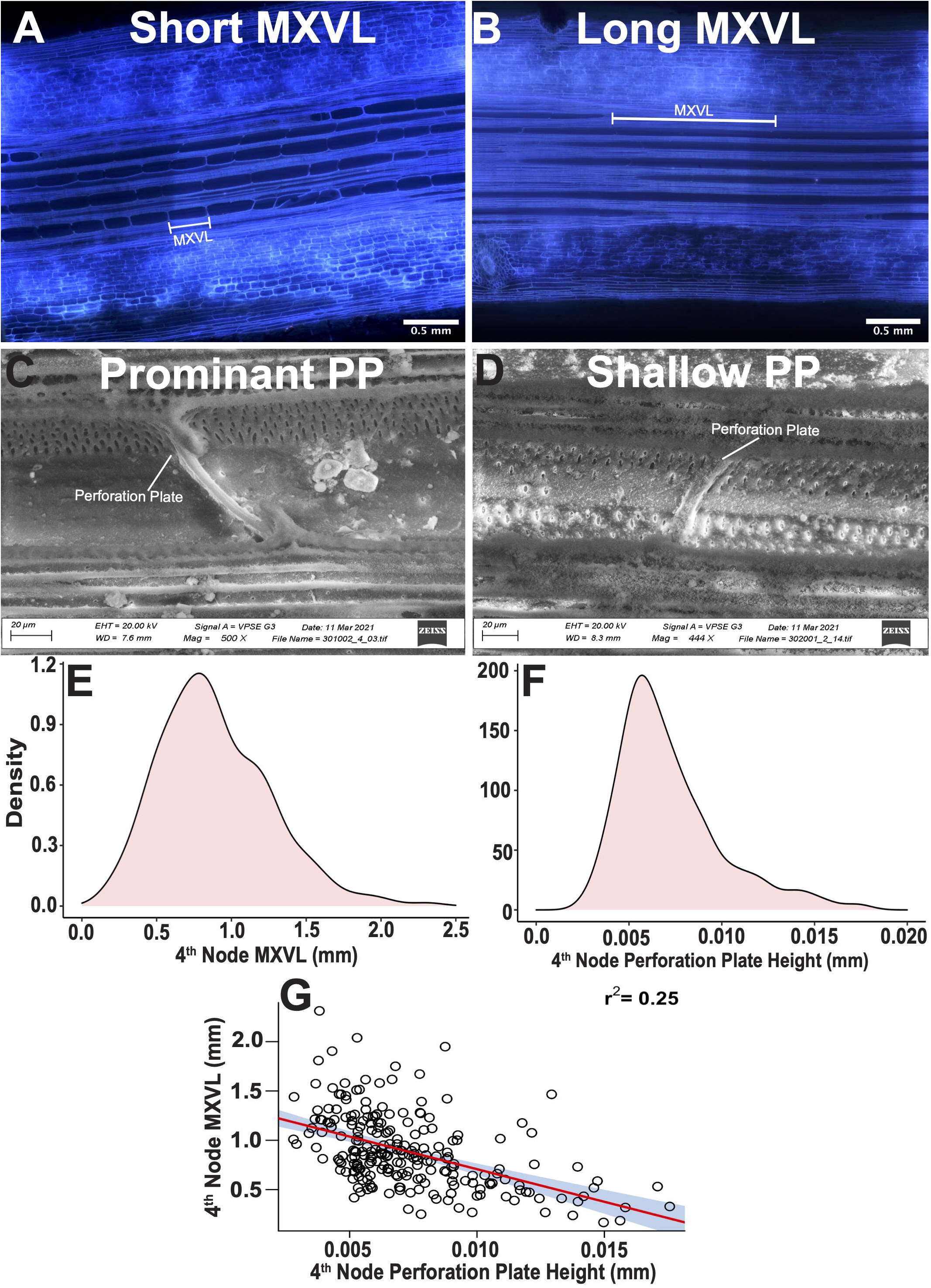
Longitudinal sections of maize (*Zea mays* L.) roots showing variation for perforation plates, density plots show the distribution of metaxylem vessel length and perforation plate height, and a regression between measurements of perforation plate height and metaxylem vessel length.

### Quantitative Trait Mapping

QTL mapping for perforation plate height and MXVL was performed using samples phenotyped from the IBM population (B73 x Mo17) grown at the Ukulima Root Biology Center described above. A subset of 141 genotypes from the IBM population for which we had phenotypic data and genotypic data for 1106 SNP markers were used for this work. Composite interval mapping was used to identify QTL with five marker covariates and a window size of 10 cM in R/qtl (Broman *et al*., 2003). The LOD threshold was determined using 1000 permutations at a significance threshold of 0.05. The Haley-Knott regression method was used to refine the positive effect of significant QTL. Marker number, length, and average and maximum spacing of the genetic map has been previously published. The physical position of markers is based on the version 4 B73 (AGPv4) reference sequence assembly (Jiao *et al*., 2017). Further details regarding the IBM population and genetic maps are provided in Burton *et al*., 2014.

### Genome-Wide Association Mapping

Perforation plate height and MXVL were phenotyped using LAT in 469 genotypes of the Wisconsin Diversity Panel. The Wisconsin Diversity Panel is composed of inbred lines that display uniform vigor and reach physiological grain maturity in the Northern Midwest of the United States. Plants were grown at the Apache Root Biology Center in Wilcox, AZ, USA (32.032079°N, 109.691171°W) in 2016 and samples from forth node roots were collected for anatomical analysis at anthesis. Averages over reps of perforation plate height and MXVL were used for further analysis. BLINK (Bayesian-information and Linkage-disequilibrium Iteratively Nested Keyway; Huang *et al*., 2019) method implemented in R package GAPIT was found to be the best fit for both perforation plate and MXVL. Significant SNPs were identified based on a genome-wide corrected Bonferroni threshold of -log(p) = 7. Data analysis was performed using R software and Bioconductor (Gentleman *et al.,* 2004). MapMan (Usadel *et al.,* 2009), and MaizeGDB (Lawrence, 2005) were used to annotate genes. Significant SNPs identified in GWAS were translated to candidate genes based on the physical position of the genes in the version 4 B73 (AGPv4) reference sequence assembly (Jiao *et al.,* 2017).

### Computational Fluid Dynamics Models

A three-dimensional model of metaxylem vessels was generated in SolidWorks based on the range of measures observed for vessel and perforation plate dimensions in the IBM population. All models were parameterized with a conduit length 3.0 mm and a vessel diameter of 0.095 mm, which was the median vessel diameter observed in the IBM population. Distance between sets of plates along the length of the conduit (MXVL) was parameterized to either 1.00 mm or 0.5 mm, reflecting the average range of MXVL in IBM genotypes with long or short vessels, respectively. Perforation plate height was parameterized to restrict the xylem lumen down to 80% of the xylem lumen (0.005 mm plate height in a 0.095 mm diameter conduit) or down to 10% of the xylem lumen (0.033 mm plate height in a 0.095 mm diameter conduit) as was observed in samples from the IBM population. The angle of the perforation plates was perpendicular (90°) to the lumen wall and the width of perforation plates was parameterized to 8 μm in all models. Flow resistance of each model was determined by the difference in pressure between the model inlet and outlet to maintain a flow rate of 2.088×10^-11^ m^3^ s^-1^. PowerCube-S01 with a high-performance computing system was used for the simulation.

### Root Elongation Rate Study

Seeds of IBM genotypes with contrasting MXVL phenotypes (IBM111 and IBM177) were germinated in root boxes in a growth chamber for 48 h, then moved under growth lights. Four replicates of each genotype were grown. To determine elongation rates the primary root of each seedling was imaged every day for 5 d. Changes in root length over the 5 d period were measured using ImageJ (Schneider *et al*., 2012).

### Vulnerability to Cavitation Study

Vulnerability to cavitation was measured using a modified version of the optical methods described by Gauthy *et al*., 2020. Three IBM genotypes with contrasting MXVL phenotypes were grown in Hagerstown-Opequon series soil at Mr. Toad’s Glee Club and Research Farm (Boalsburg, PA; 40.794655°N, -77.764351°W). Ten cm of the fifth leaf apex was harvested once fully mature and hydrated in wet paper towels for 2 h before imaging. Leaf segments were affixed to a diffuse LED light source with adhesive tape heated to 37°C while imaging a region of interest 5 cm from the cut leaf edge every 1 min on a Nikon SMZ 1500 stereo microscope. Images were converted to 8-bit grayscale before subtraction of sequential frames in ImageJ to visualize cavitation events by optical refraction/darkening. Due to the parallel venation of maize leaves, medial shrinkage/slippage caused artifacts that prevented automated analysis of cavitation. ImageJ was used to manually measure the length of cavitation events on each frame for comparison between genotypes.

### Controlled Environment Study

This study was conducted in a glasshouse located at Pennsylvania State University in University Park (40.801955°N, 77.862544°W). Plants were grown from April through June 2021 under a 16/8-h (light/dark) photoperiod, 40% relative humidity, and maximum/minimum temperatures of 28°C/26°C. Midday photosynthetic active radiation was 900 to 1,000 μmol photons m^-2^s^-1^. Natural light was supplemented from 06:00 to 22:00 with approximately 500 μmol photons m^-2^s^-1^ from metal-halide lamps. Seeds were surface sterilized in a 25% (v/v) NaOCl in water for 2 min, rinsed in deionized water, and germinated in 0.5 mM CaSO_4 i_n the dark at 28°C for 72 h. Uniform seedlings were transplanted to the glasshouse in opaque, 30 L mesocosms 15 cm in diameter and 155 cm in height and lined with transparent 6 mm high-density polyethylene film to facilitate root sampling. Mesocosms were filled with a mixture of 4% (w/w) coarse grade A perlite (Whittemore), 50% (w/w) medium-grade sand (US Silica), 26% (w/w) D3 coarse grade A vermiculite (Whittemore), and 20% (w/w) field soil (Ap2 Hagerstown silt loam [fine, mixed, semiactive, mesic Typic Hapludalf]) sieved through 6 mm mesh. The soil was incorporated to replicate features found under field conditions, such as the presence of organic matter, soil biota, and oxide surfaces that serve to buffer nutrient availability. Mesocosms were fertilized with 5 g kg^-1^ Osmocote (15-9-12; 5-6 mo.) (The Scotts Co., Marysville, OH) incorporated into the media at the time of mixing and consisting of (%): NO_3 (_8) NH_4 (_7), P (9), K (12), S (2.3), B (0.02) Cu (0.05), Fe (0.68), Mn (0.06), Mo (0.02), and Zn (0.05).

A Randomized Complete Block Design was utilized with two irrigation levels; water stress (WS) and well-watered (WW). Irrigation was supplied through drip rings, with pots being brought to field capacity daily. Irrigation was halted on pots assigned to the WS treatment at 17 days after planting (DAP). The experiment was run for a total of 42 d with destructive measurements taken from all genotypes in all treatments at 17, 31, and 42 DAP. Each genotype had four replications at each time point and treatment.

To determine net water loss from pots in the WS treatment, 5 pots from each phenotypic group were weighed hourly from 17 to 42 DAP using Adam CPWplus 75 floor scales (Adam Equipment, Oxford, CT). Gravimetric soil moisture was also determined at 31 and 42 DAP at 20 cm increments by depth from the soil surface to the bottom of the container. Leaf relative water content and specific leaf area were determined from 5, 2.5 cm leaf discs collected at 16:00 on 41 DAP as in Smart and Bingham (1974). Predawn water potential of plants in the WS treatment was determined using the PMS model 615 Scholander pressure bomb at 42 DAP from 03:00 to 06:00 (PMS Instruments, Albany, OR).

At 17, 31, and 42 DAP destructive measurements were taken including leaf number, leaf area, internode distance, plant height, as well as dry root and shoot biomass. Leaf area was determined using the Li-Cor LI-3100C leaf area meter (Li-Cor, Lincoln, NE). Dry mass was determined from tissues dried at 65°C for 7 d.

At 31 DAP, two, 10 cm segments of nodal roots from the most recently emerging node were collected 5 cm from the base of the root for measurement of root respiration. Root respiration rates were determined immediately after excavation and washing using a Li-Cor 6400 gas-exchange system with a modified respiration chamber (Li-Cor, Lincoln, NE). Lateral roots were removed from these segments with a razor prior to respiration measurements. Measurements were performed under ambient glasshouse conditions, with the sealed chamber being kept at a temperature of 28°C and baseline sample chamber and reference chamber CO_2 c_oncentration of 400 μmol mol^-1^.

42 DAP, mesocosms were cut into 20 cm increments by depth from the soil surface to the bottom of the container. Roots were washed, collected, and imaged from each 20 cm segment using an EPSON Perfection V700 PHOTO scanner and total length was quantified with WinRhizo software (WinRhizo Pro; Reagent Instruments). The scanned roots were then dried and weighed to determine specific root length, calculated by dividing the total root length by the total root dry weight.

At 42 DAP, two, 10 cm segments of nodal roots from the most recently emerging node were collected 5 cm from the base of the root. Lateral roots were then removed with a razor and *in situ* measures of axial conductance were then performed on these root segments as in Strock *et al*., (2021). Prior to measurement, each 10 cm root segment was soaked in a de-gassed 20 mM KCl solution for 30 min. Paraffin wax was melted and painted on the surface of the root to preclude radial losses of flow through lateral root junctions across the segment. A 0.0093 MPa hydraulic head of degassed 20 mM KCl solution was attached to one end of the root segment and flow out the opposite end of the segment was quantified over a 1 min period using an Adventurer Pro AV13C analytical balance (Ohaus Corporation, Pine Brook, NJ). Following measurement of axial conductance across the 10 cm segment, 2.5 cm increments of root were subsequently excised from the end of the segment and the measurement was repeated with 7.5, 5, and 2.5 cm lengths of root to determine the effect of perforation plated on axial flow.

Following these *in situ* conductance measurements, the wax coating was removed, and the root segments were preserved in 75% (v/v) ethanol in water. Preserved segments were sectioned in both the longitudinal and cross-sectional dimensions with laser ablation tomography (Strock *et al*., 2019). Length of metaxylem vessels was measured in the longitudinal dimensions while the number and area of metaxylem vessels was measured from the cross-sectional dimension at both ends of the segment. Theoretical axial metaxylem conductance (k_h;_ kg m MPa^-1^ s^-1^) was calculated for each cross-sectional image using the modified Hagen-Poiseuille law (Eq. 1), where d is the diameter of the vessel in meters, ρ is the fluid density (equal to water at 20°C; 1000 kg m^-3^), and η is the viscosity of the fluid (equal to water at 20°C; 1 × 10^-9^ MPs s^-1^; Tyree and Ewers, 1991). The mean theoretical conductance estimate calculated from images at each end of the segment were used for comparison with the *in situ* measure of conductance across that segment.

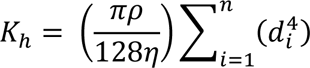

**Equation 1**. Modified Hagen-Poiseuille law.

### USA Field Trials

The PA19 and PA20 field trials was conducted at the Russell E. Larson Agricultural Research Farm at Rock Springs, PA, USA (40.711365°N, 77.953089°W) from June through September 2019 and 2020, respectively. The soil at this site is a Hagerstown silt loam (fine, mixed, mesic Typic Hapludalf). A split plot design was utilized with two irrigation levels; two, 0.02 ha rainout shelters were split into two, 0.01 ha blocks each, and two 0.02 ha irrigated fields split into two, 0.01 ha blocks. In 2019, sixteen genotypes and in 2020, four genotypes were randomized within each block. To manage fungal pathogens, seed were treated with Captan 50W fungicide solution (0.2g/ L) at a rate of 0.5 ml/100 seeds prior to planting. All fields were fertilized to meet the nutrient requirements of maize as determined by soil tests at the beginning of the season. Each genotype was planted in a single row, 4.6 m long plot with 76 cm row spacing at a density of 73,300 plants ha^-1^. During periods of inadequate rainfall, irrigation was supplied to the well-watered treatment. Drought treatment was initiated at 20 DAP, after which water-stressed plots experienced no rainfall or irrigation through the time of yield harvest. Each genotype had four replications within each irrigation treatment.

At anthesis, leaf, brace root, and forth node root samples were collected for anatomical analysis as described above. Dry masses were determined from tissues dried at 60°C for 7 d.

### Graneros, Chile Field Trial

The CL20 field trial was conducted at the Tuniche Research Farm near Graneros, Chile (-34.108279°S, -70.748495°W, soil order is Inceptisol) from November 2019 through May 2020. A split plot design was utilized with two irrigation levels; one field where irrigation was limited was split into two water-stressed blocks, and one irrigated field split into two blocks. Thirty hybrid genotypes known to contrast in water use efficiency (15 drought tolerant genotypes, 15 drought sensitive genotypes) were randomized within each block. All fields were fertilized to meet the nutrient requirements of maize. Each genotype was planted in a two row, 4.6 m long plot with 76 cm row spacing. During periods of inadequate rainfall, irrigation was supplied to the well-watered treatment. Each genotype had four replications within each irrigation treatment.

At anthesis, destructive measurements were taken including dry shoot biomass, leaf samples and forth node root samples for anatomical analysis as described above. Dry masses were determined from tissues dried at 60°C for 7 d.

### Statistical Analysis

All statistical analyses were performed in R v. 3.6.2 (R Core Team, 2019) with graphical illustration using “ggplot2” (Wickham, 2016). Prior to all statistical tests, the normality and homoscedasticity of the data were determined using the Shapiro-Wilk test and the nonconstant error variance test, respectively. Where data did not meet these assumptions, a box-cox or log transformation was used to help normalize the data. Significant correlations and differences for all data analyses were considered at α ≤ 0.05 and at α ≤ 0.1 where noted.

Broad-sense heritability (H^2^) and repeatability (R^2^) were calculated for MXVL in below-ground 4^th^ node roots, above-ground brace roots, and leaves on an entry-mean basis according to Fehr (1987) where σ^2^(G) is the genotypic variance and σ^2^(E) is the error variance (Eq. 2). Genotypic and error variance were generated for each phenotype from linear mixed models fit by the restricted maximum likelihood (REML) method. In the determination of variance components, each occasion that a genotype was germinated in was considered a replicate.

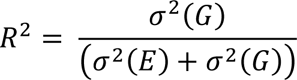

**Equation 2**. Calculation of repeatability (R^2^)

## Results

### Perforation plate phenotypes vary among maize genotypes

Considerable variation was observed among maize genotypes for the height (prominence) of perforation plates extending into the xylem lumen as well as the frequency of perforation plates occurring along the length of metaxylem conduits (MXVL) (Fig. 2A-2D). The distribution of these phenotypes in the IBM population was close to normal or slightly skewed to smaller values (Fig. 2E, 2F). A significant relationship between the height of perforation plates and length of metaxylem vessels was observed (r^2^=0.25) where genotypes with short MXVL tended to have taller perforation plates while genotypes with long MXVL tended to have shorter, less prominent perforation plates (Fig. 2G).

IBM genotypes with contrasting MXVL phenotypes in nodal roots displayed similar differences in MXVL of above-ground brace roots and leaf tissue (Fig. 3, S1). Water stress did not have a significant effect on MXVL phenotypes (Fig. 3, S1). No other differences in metaxylem phenotypes such as the number of metaxylem vessels, the total cross-sectional area of metaxylem vessels, or the estimated conductance of root segments were observed among genotypes with contrasting MXVL phenotypes (Supplemental Fig. S2). Genetic factors explained 77.7%, 61.8%, 43.3% of phenotypic variation for MXVL in nodal roots, brace roots, and leaves respectively (Table 1). Similarly, genetic factors explained 37.8%, 49.0%, 28.6% of phenotypic variation for perforation plate height in nodal roots, brace roots, and leaves respectively (Table 1).

**Figure 3.**
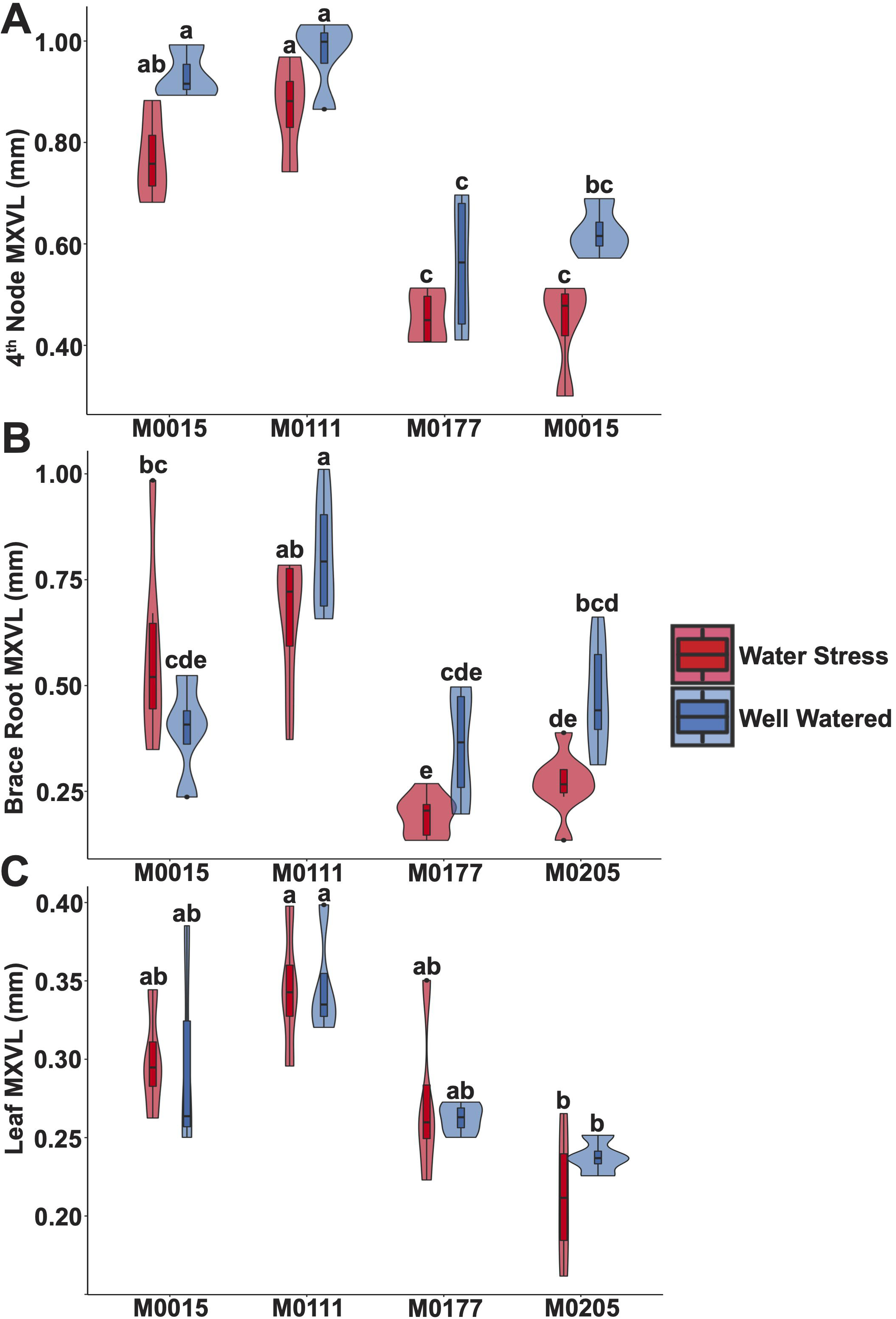
Violin plots of MXVL in below-ground 4^th^ node roots, above-ground brace roots, and leaves in the four IBM genotypes of maize (*Zea mays* L.) under water stress and well-watered conditions.

**Table 1.**
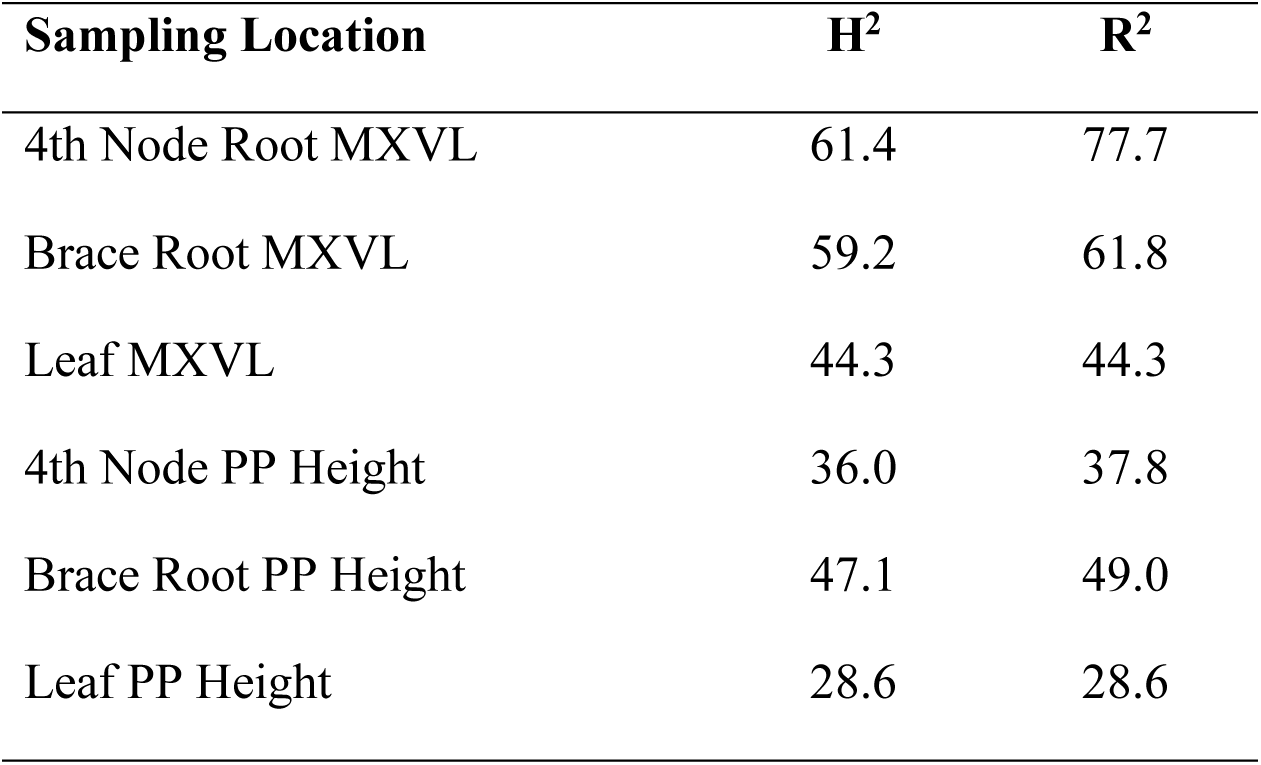
Broad-sense heritability (H^2^) and repeatability (R^2^) of MXVL and PP Height in the four IBM RILs of interest in below-ground axial roots from node 4, aboveground brace roots and leaves. Heritability and repeatability were calculated for 4^th^ node roots measured at field sites in South Africa, the United States and Chile under water deficit and well-watered conditions and calculated for brace roots and leaves in the United States and Chile sites under water deficit and well-watered conditions.

### Genetic control of perforation plates in maize

Within the IBM population, QTL were detected for both MXVL (Fig. 4A) and perforation plate height (Fig. 4B). For MXVL, QTL were identified on chromosome 2, explaining 9.13 % of the phenotypic variation. For perforation plate height, QTL were identified on chromosome 3, explaining 10.29 % of the phenotypic variation for this phenotype.

**Figure 4.**
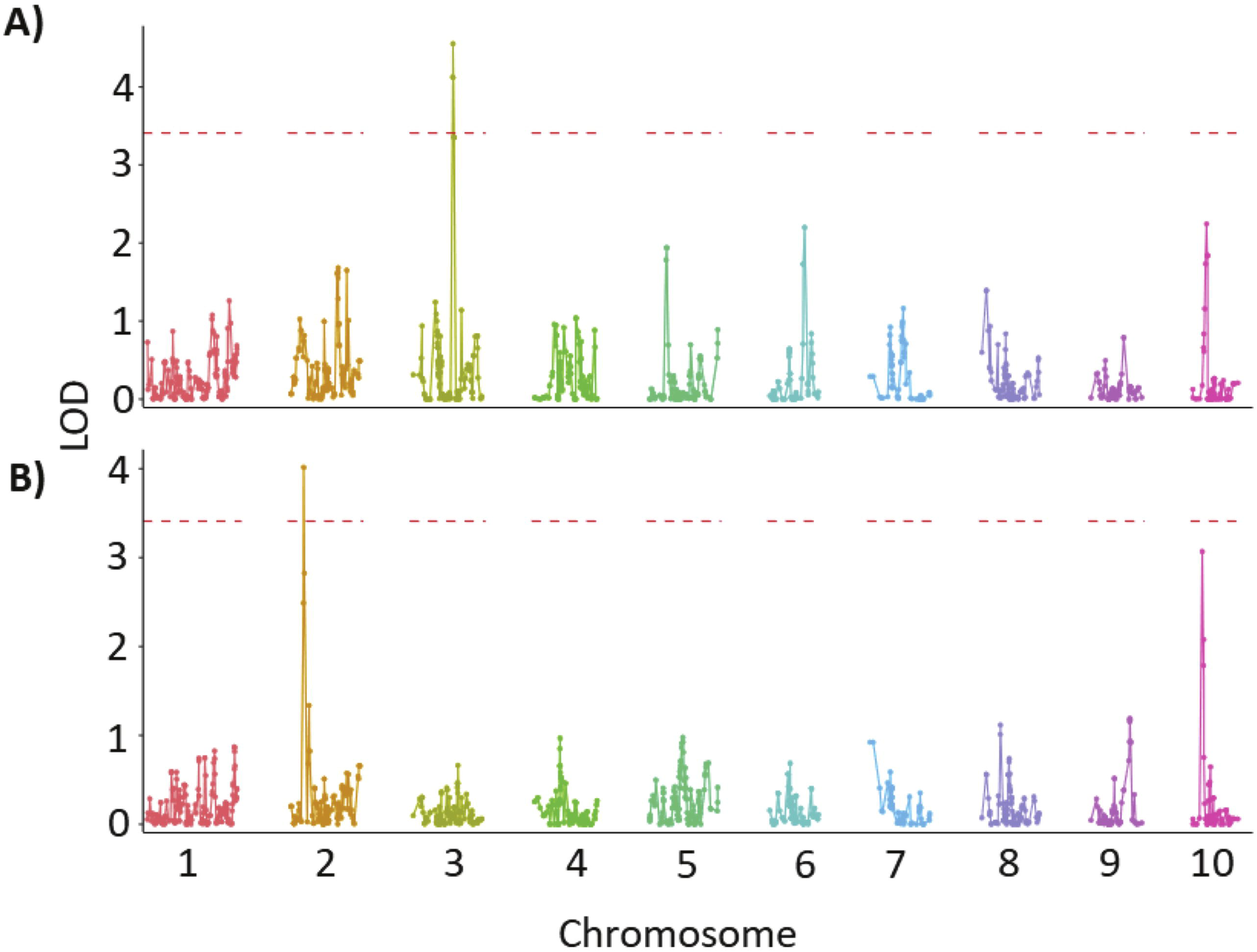
Composite interval mapping for MXVL and plate height show genetic control of these features in the biparental population (B73 x Mo17) of maize (*Zea mays* L.).

Genome wide association mapping/studies (GWAS) on the Wisconsin Diversity Panel identified two significant SNPs for MXVL using a Bonferroni-corrected genome-wide threshold value of - log(p) = 7. Candidate genes were selected from gene models containing SNPs above the Bonferroni significance threshold. One of the significant SNPs was located in gene model ZM00001d030760 on chromosome 1, which is predicted to be a Asparagine synthase family protein that is expressed primarily in roots under drought stress (Fig. 5A). The other significant SNP for MXVL was linked to the Zm00001d012015 gene model on chromosome 8 predicted to be a Squamosa promoter-binding 2 protein that is largely expressed in roots (Fig 5A). Two significant SNPs for plate height were identified, one on chromosome 1 and another on chromosome 8 (Fig 5B). Four candidate genes were found linked with SNP on chromosome 1 including Zm00001d031943 (Kinesin heavy chain), Zm00001d031944 (Hairpin-induced protein), Zm00001d031945 (Glutathione S-transferase, N-terminal domain containing protein), and Zm00001d031946 (unknown function). Chromosome 8 SNP was found in Zm00001d011745 which belongs to Lecithin:cholesterol acyltransferase family.

**Figure 5.**
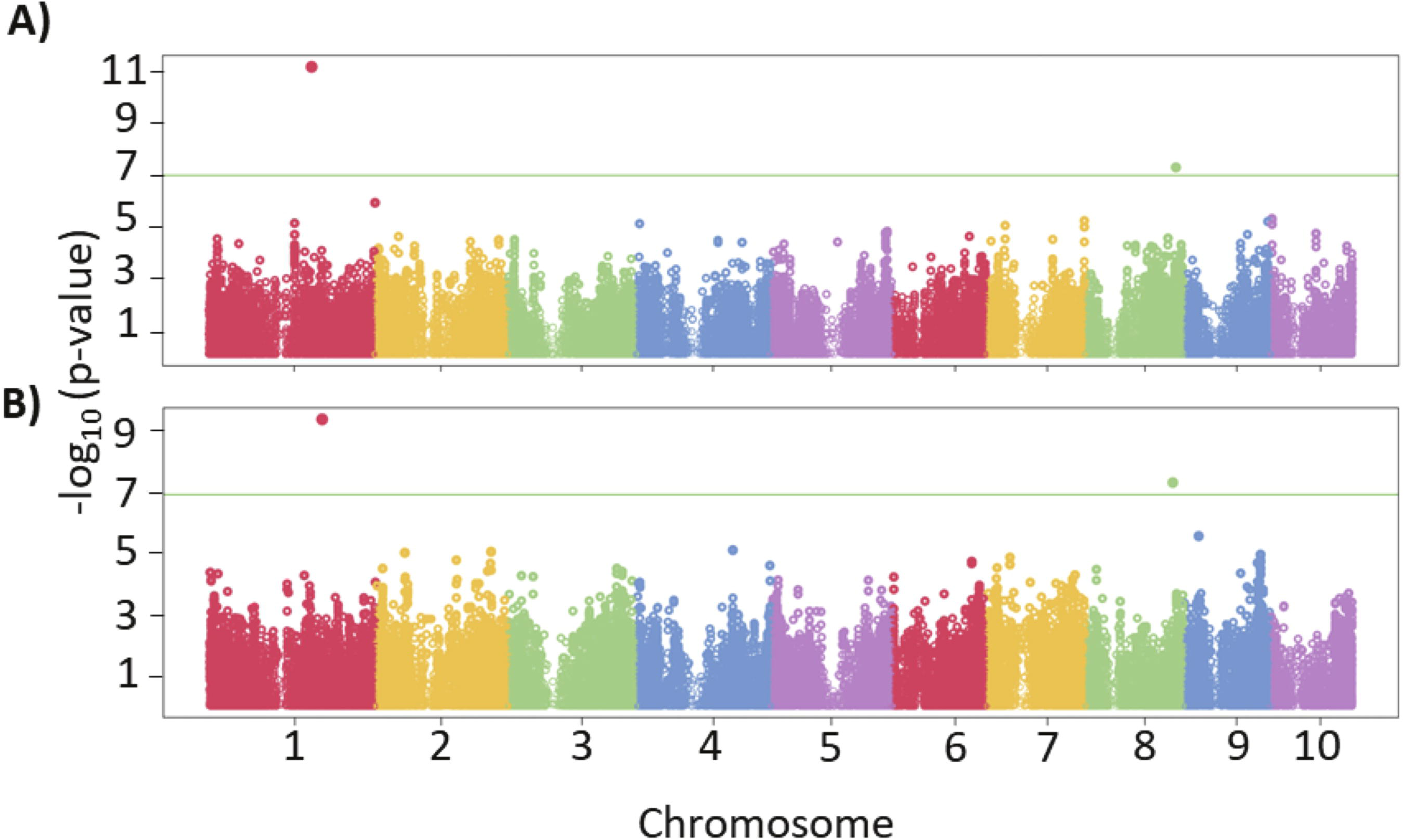
Manhattan plots for MXVL and plate height showing genetic control for these features in the Wisconsin Diversity Panel of maize (*Zea mays* L.).

### Effect of perforation plate phenotypes on water transport

Computational fluid modeling suggested that over a 3.0 mm long vessel, perforation plate height has a much larger influence on flow resistance than does MXVL (Table 2). In actual root segments, the difference between conductance estimated from the number and diameter of vessels using the Hagen-Poiseuille law and the actual conductance measured across the root was well described by the length of metaxylem vessels in that root segment (r^2^ = 0.35) (Fig. 6B). Additionally, the influence of perforation plates on actual conductance is reflected in the increased conductance of the root segment as its length is reduced (Fig. 6C).

**Figure 6.**
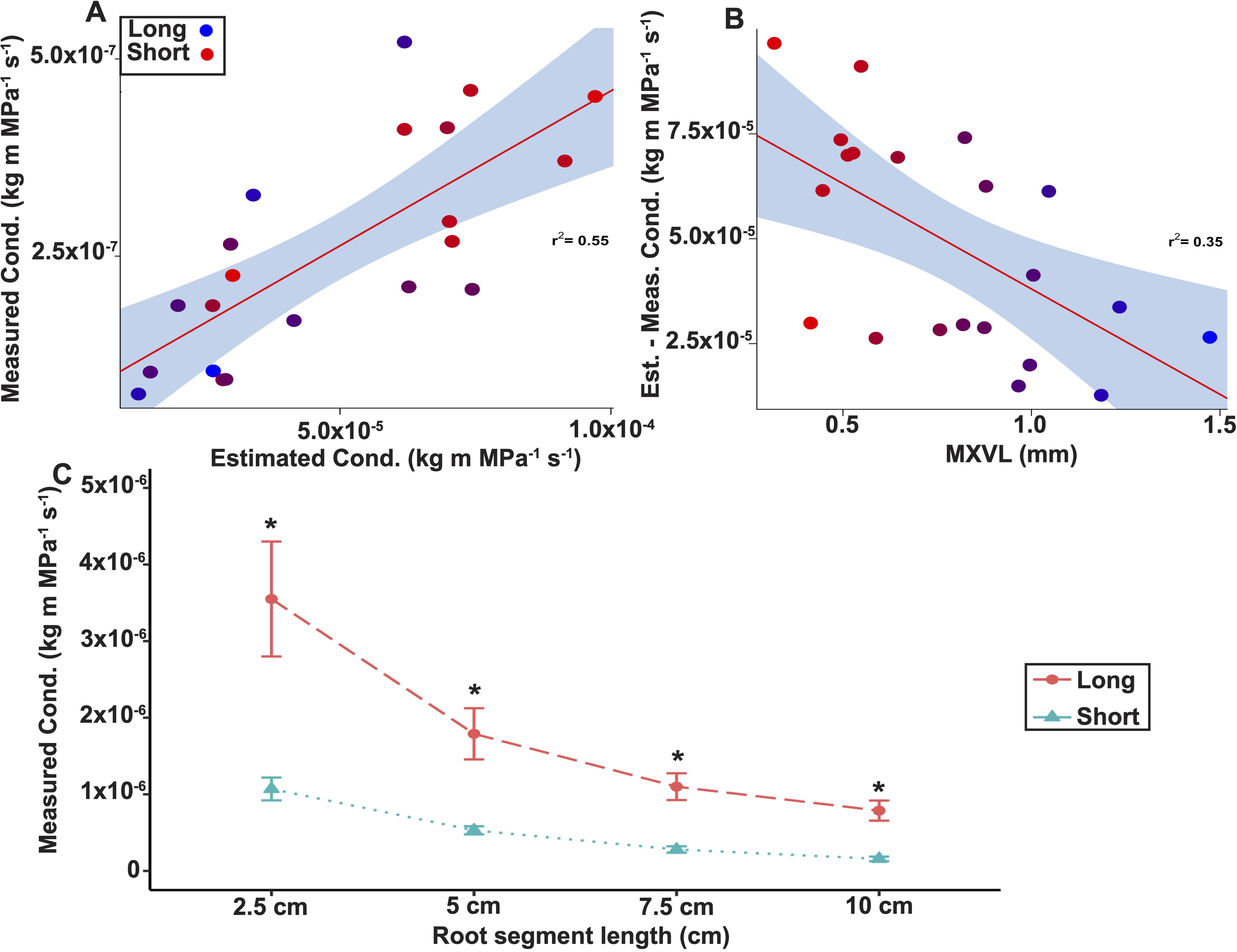
The relationship between axial conductance *in situ*, conductance predicted by the Hagen-Poiseuille law, and MXVL of field grown root segments.

**Table 2.**
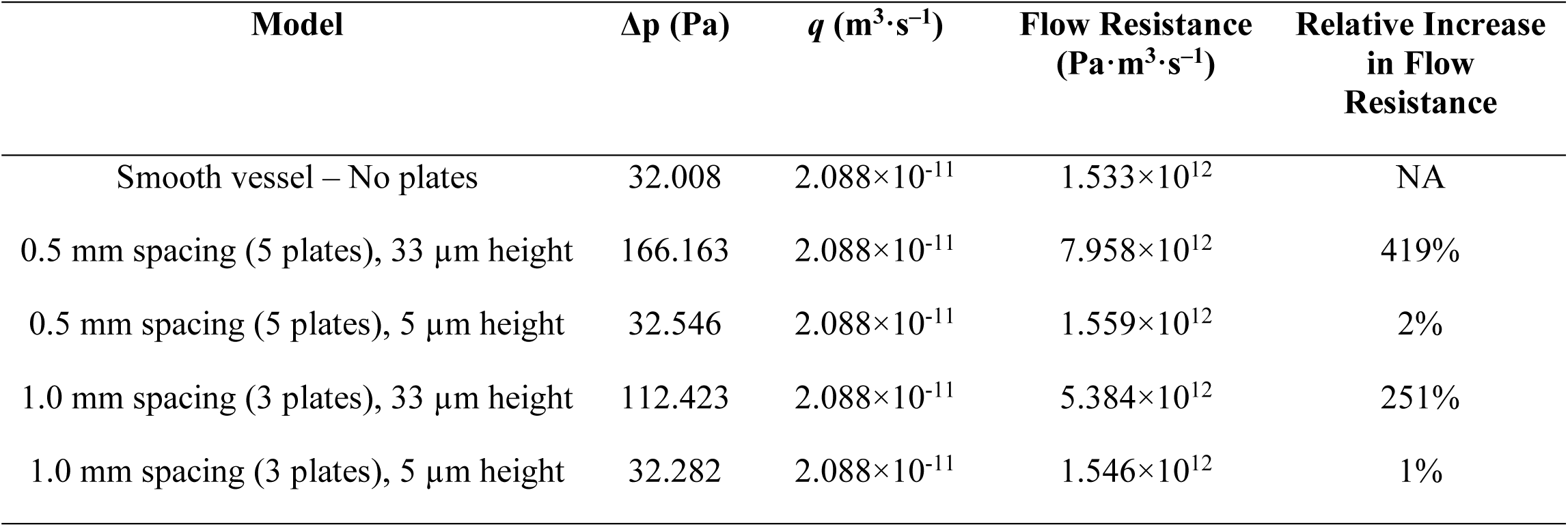
The pressure differential (Δp), average flow rate (*q*), flow resistance (Δp/*q)*, and the increase in flow resistance relative to a smooth vessel with no perforation plates in computational fluid velocity models of a 3.0 mm vessel with different combinations of perforation plate height and spacing.

### Effect of perforation plates on vulnerability to cavitation

No clear differences in vulnerability to cavitation were observed among genotypes with contrasting perforation plate phenotypes. Initial root cavitation measurements were obscured by the optical effects of drying root mucilage. Because MXVL phenotypes were pleiotropic between roots and leaves (Fig. 3), all cavitation experiments were performed on excised leaf segments for image clarity. Leaves grown in the growth chamber were thin and soft, and did not display any measurable cavitation events. Mature leaves grown in the field had variable cavitation that occurred over a period of approximately 2-5 h (Supplemental Fig. S3A). To compare cavitation between leaves, normalized data was used to identify a threshold of 50% cavitation (Supplemental Fig. S3B). There was no significant difference in the 50% cavitation threshold among metaxylem vessel length phenotypes (Supplemental Fig. S3C).

### MXVL is associated with root elongation

MXVL phenotypes are linked to the elongation of different plant organs. Genotypes with the long MXVL phenotype displayed greater rates of root elongation (Fig. 7A, 7B) and greater total root length by 42 DAP under water deficit (Fig. 7C). Despite these differences in root elongation rates, no differences in root respiration rates between long or short MXVL phenotypes were detectable at either apical or basal root segments (Supplemental Fig. S4).

**Figure 7.**
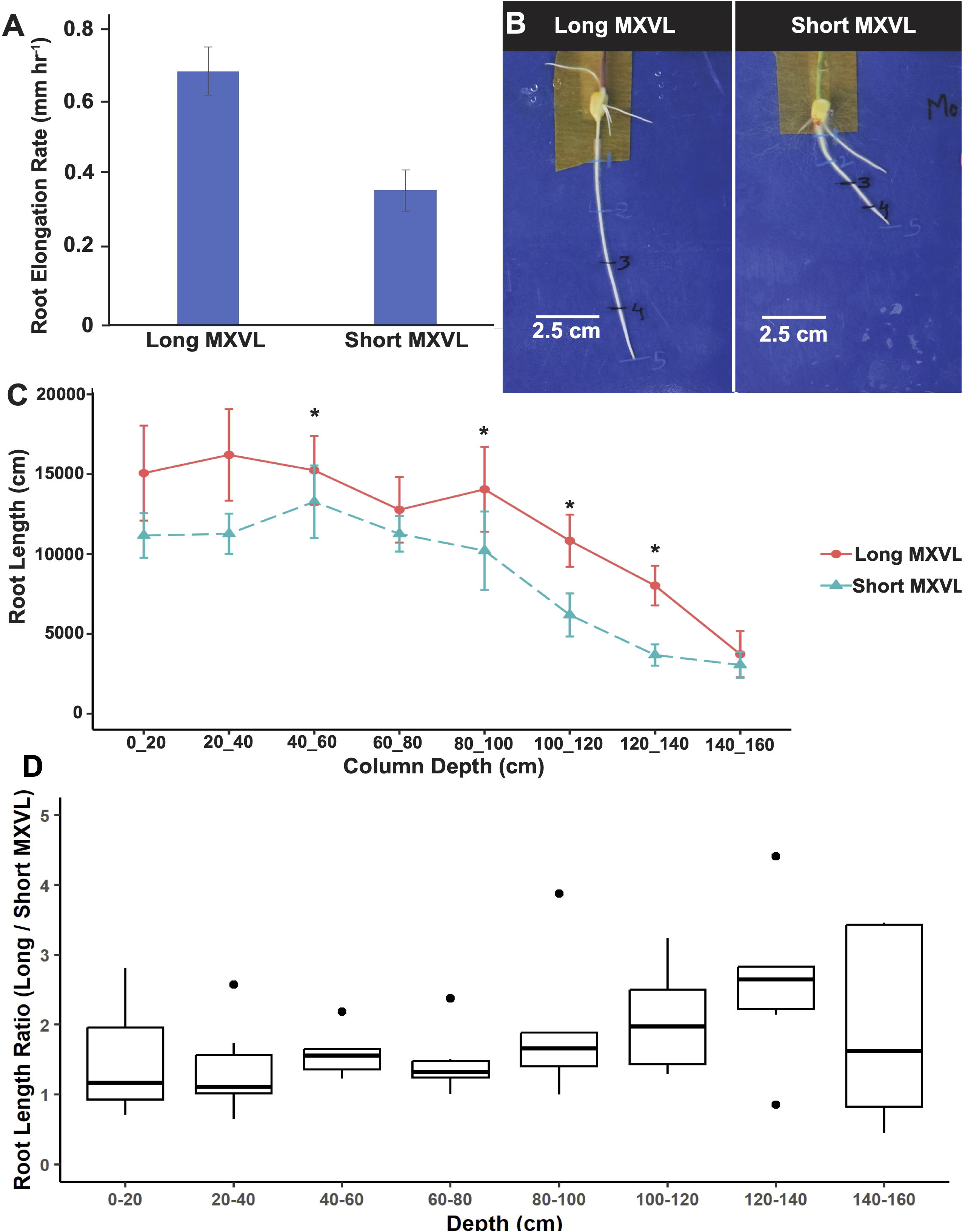
Root elongation rates measured in the primary roots of maize (*Zea mays* L.) seedlings, root length distribution in the glasshouse at 42 DAP under drought stress, and the ratio of total root length (long/short MXVL) across soil depth.

Genotypes with the long MXVL phenotype also tended to have taller shoots with longer internode distances than genotypes with the short MXVL phenotype (Supplemental Fig. S5B, S5C). No differences were observed in shoot mass, leaf area, or specific leaf area in the glasshouse between long and short MXVL genotypes.

### Relationship of MXVL with plant water status under water deficit

A significant effect of MXVL phenotype on gravimetric soil water content sampled across the depth of the mesocosm was observed at 31 DAP (Fig. 8B; p = 0.049) and 42 DAP (Fig. 8C; p = 0.022) , with the short MXVL genotype having more water at 120-140 cm depth at 42 DAP (Fig. 8C). Despite having more available water at depth, by 42 DAP under water deficit, genotypes with the short MXVL phenotype had lower predawn leaf water potential (p = 0.07) and midday leaf relative water content (p = 0.07) than genotypes with the long MXVL phenotype (Fig. 9).

**Figure 8.**
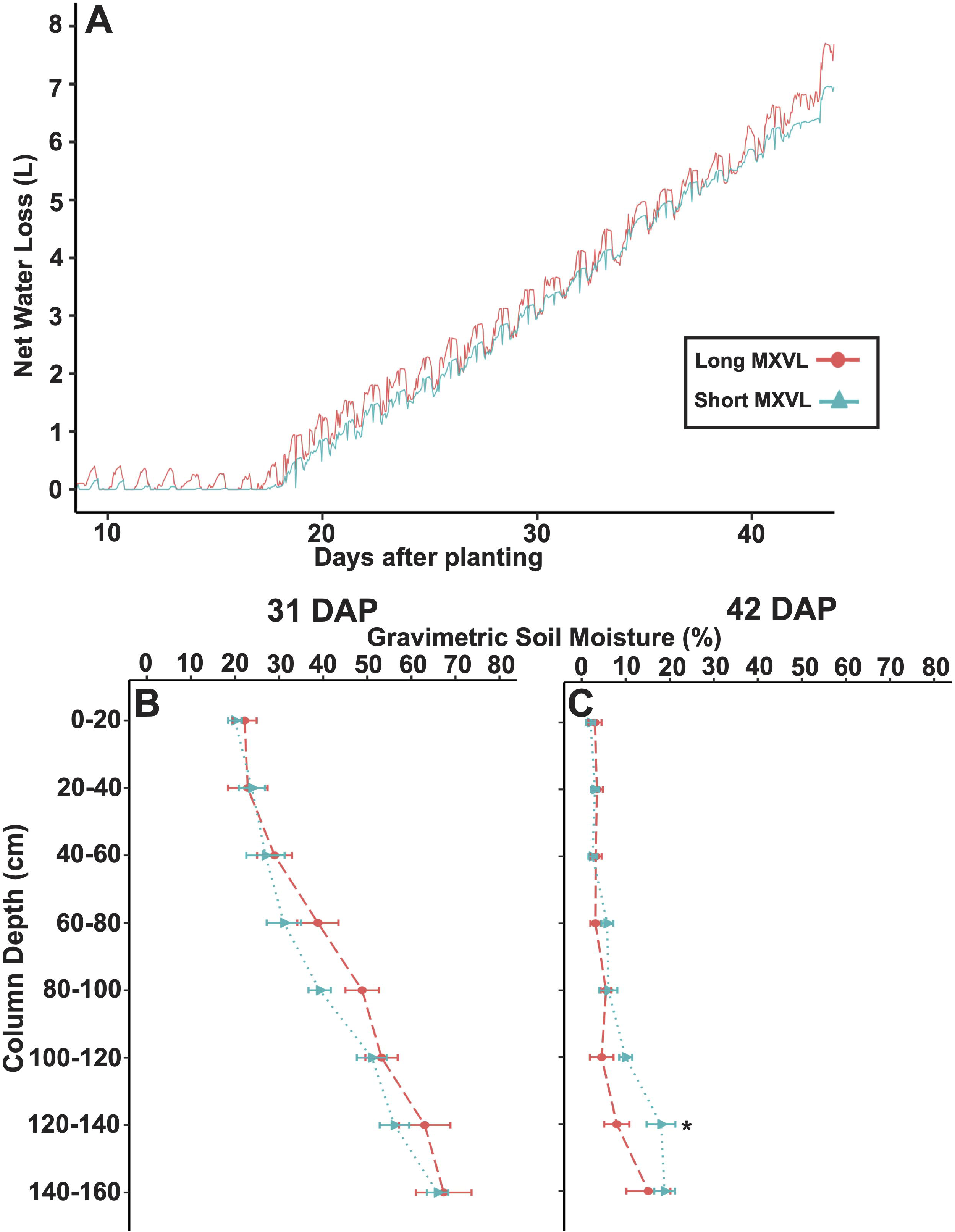
Water loss (L) from mesocosms planted with maize (*Zea mays* L.) in the glasshouse and gravimetric soil water content (%) for mesocosms in the water deficit treatment at 31 DAP and 42 DAP.

**Figure 9.**
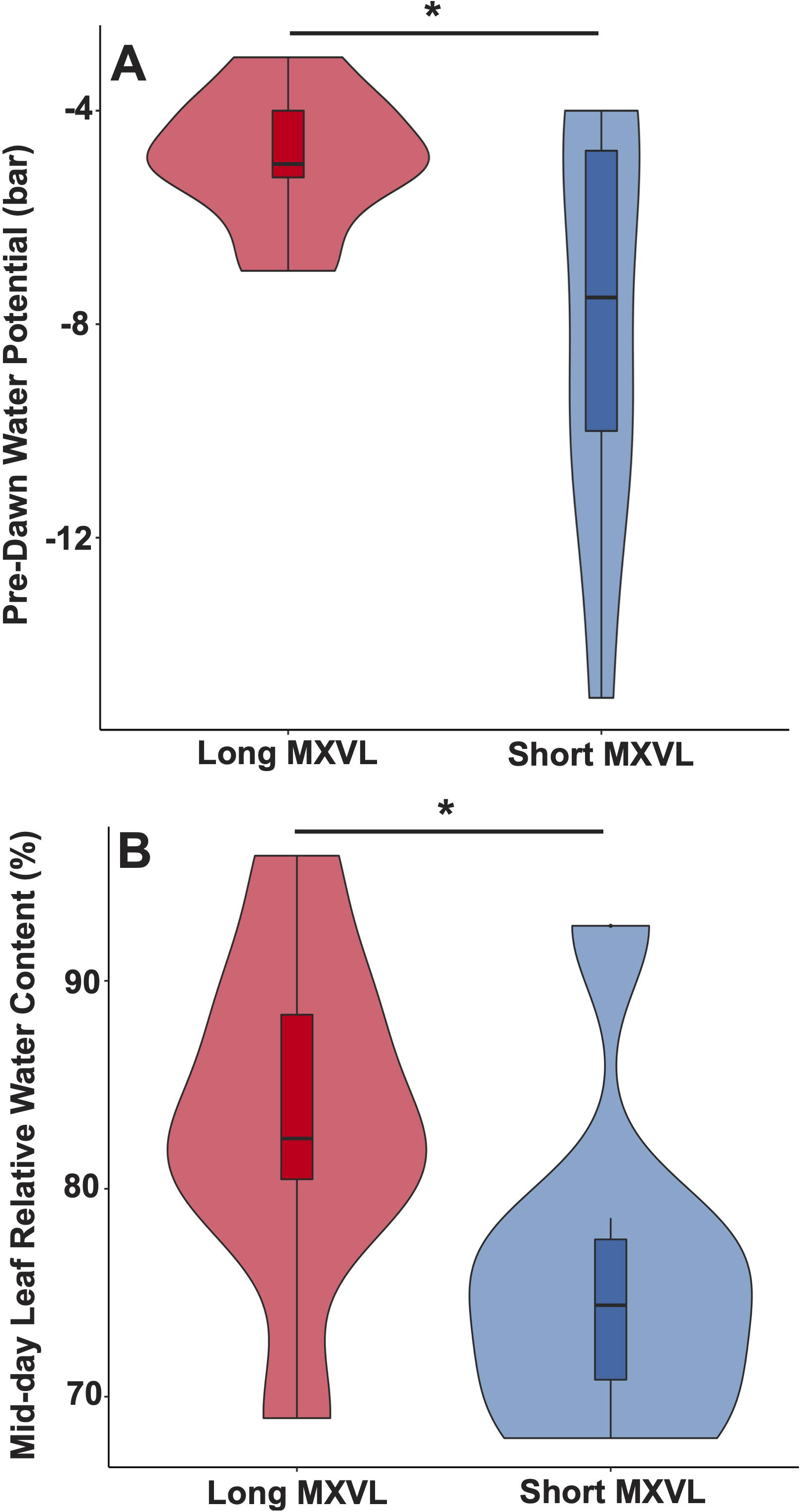
At 42 DAP under drought stress, maize (*Zea mays* L.) genotypes with the short MXVL phenotype have lower predawn leaf water potential (p = 0.07) and midday leaf relative water content (p = 0.07) than genotypes with the long MXVL phenotype.

Under terminal drought stress in the field, genotypes with the long MXVL phenotype had consistently greater yields than genotypes with the short MXVL phenotype (Fig. 10). Among a panel of 30 hybrid accessions grown in Chile, a principal component analysis of vascular phenotypes identified a primary principal component that was most significantly described by variation in root conductance, root MXVL, and distance between vascular bundles in leaves (Fig. 11C). This principal component was significantly correlated with yield, leaf rolling scores, as well as midday leaf temperature (Fig. 11B).

**Figure 10.**
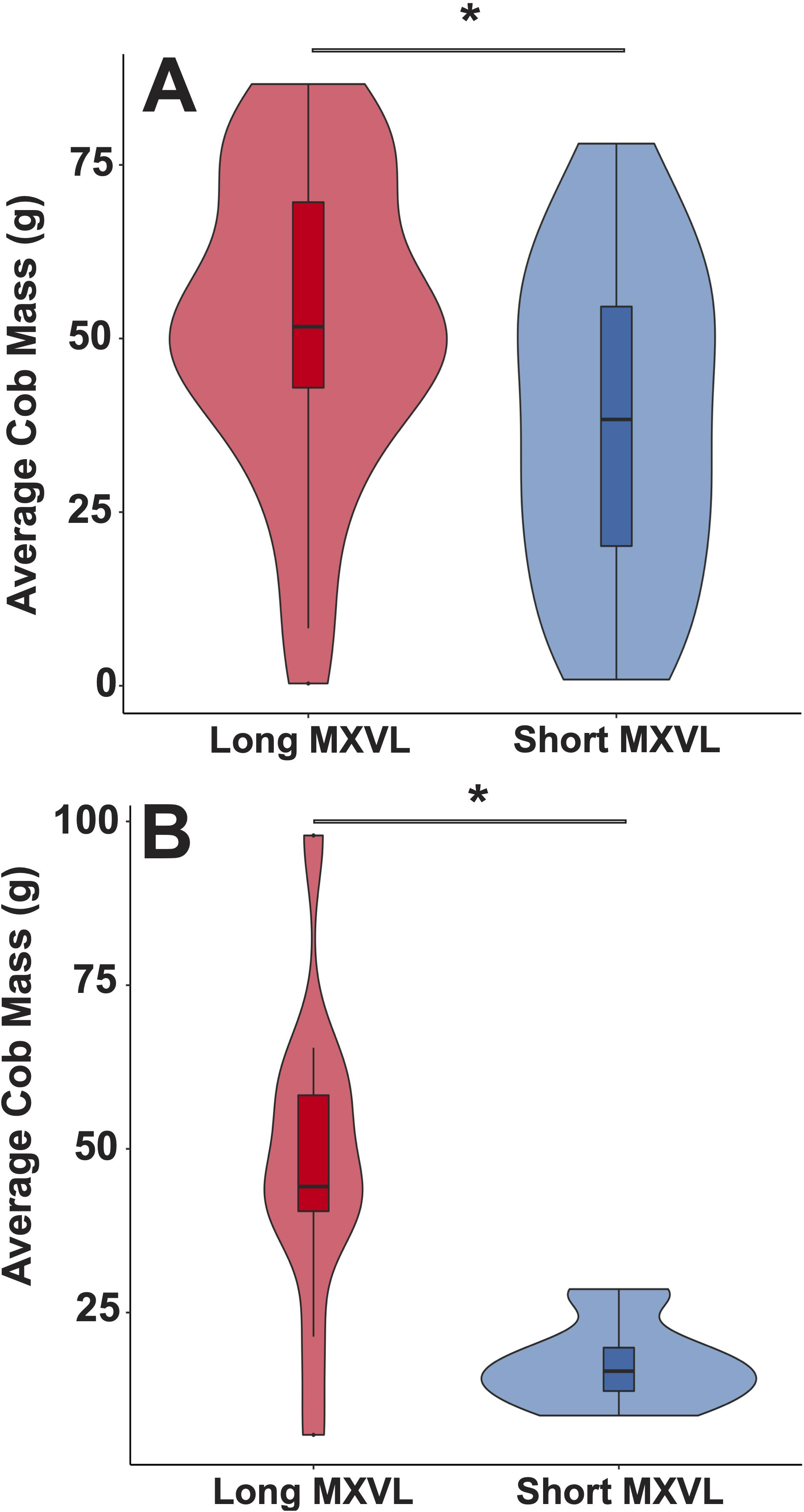
Yield of accessions from biparental population (B73 x Mo17) of maize (*Zea mays* L.) with contrasting MXVL phenotypes under terminal drought stress in the USA field trials in 2019 and 2020.

**Figure 11.**
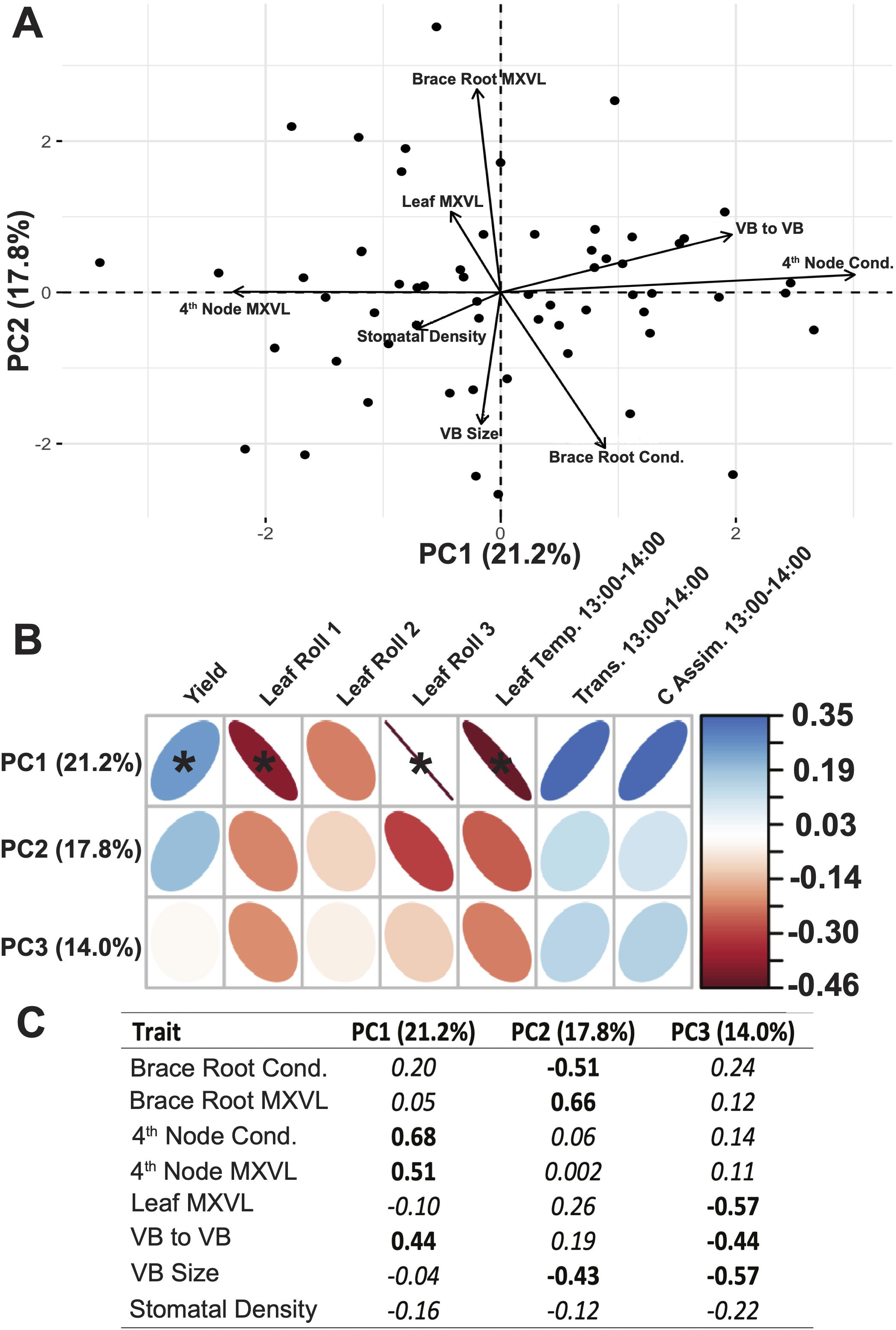
Biplot showing the relationships of root and leaf vascular phenotypes measured across 30 maize (*Zea mays* L.) hybrids grown under water stress in Graneros, Chile, as well as correlations between the first three principal components and yield, leaf rolling scores, as well as midday leaf temperature and gas exchange measures.

## Discussion

We uncovered significant intraspecific variation within *Zea mays* for the prominence and frequency of perforation plates along metaxylem vessels. Smaller plate height is highly heritable, is associated with longer vessel element length, reduced resistance to axial water transport, greater root elongation, root depth, deep water utilization, and improved plant water status under terminal drought stress. Under drought stress in the field, xylem vessel element length/perforation plate height is associated with leaf roll, leaf temperatures, transpiration, photosynthesis, and grain yield. We conclude that phenotypic variation for xylem perforation plate phenotypes in maize directly affects axial water conductance and is part of a pleiotropic syndrome with greater root elongation and deeper rooting that improves adaptation to water deficit stress. The high heritability, stability across environments, growth stages, and plant organs makes this a promising phenotype for efforts focused on improving drought stress tolerance in cereal crops.

It is noteworthy that perforation plate height and spacing directly affects axial hydraulic conductance in an annual monocot like *Zea mays*. Fluid velocity modeling suggested that the observed variation for perforation plates in maize has effects on water transport through xylem conduits, and this prediction was subsequently supported by *in situ* measures of water transport across root segments. These results align with the outcomes of modeling simulations and studies that have shown how perforation plates can play a significant role in xylem transport of perennial dicot species (Schulte & Castle, 1993a,b; Schulte, 1999; Xu *et al*., 2020, Echevarría *et al.,* 2023). A study by Xu *et al*. (2020) using a computational fluid dynamic model to explore factors influencing xylem sap flow in the dicot *Jatropha curcas,* estimated minimal hydraulic resistance of simple perforation plates to axial flow to be 3.6% of total axial resistance. Nevertheless, the range of plate heights parameterized in *Jatropha curcas* was small relative to total vessel diameter (2-5 µm in height in a 52-61 µm diameter vessel) compared to the proportions we observed in *Zea mays* (5-33 µm height in vessels 95 µm in diameter). Similar to our observations, Ellerby and Ennos (1998) show in large-scale physical models of xylem conduits that the resistance of perforation plates to axial flow is dependent upon structure and abundance along the length of the conduit. A comparative analysis across more than 1000 angiosperm species as well as finite-element modeling showed that shorter vessel elements were associated with decreased axial conductance and dryer habitats (Echevarría *et al.,* 2023). Although it was not a variable of consideration in the present work, the angle of perforation plates has also been shown to be a significant factor contributing to hydraulic resistance (Christman & Sperry, 2010).

While it is clear that genetic variation in perforation plate structure contributes to hydraulic resistance at the scale of xylem conduits, plant-level data from the glasshouse and field trials suggest that direct resistance to xylem sap flow is only one aspect of the effect of perforation plates on plant water use strategies. Considering the entire soil>plant>atmosphere continuum, simple perforation plates likely do not represent the greatest resistance to axial water transport. In maize, the points of greatest resistance within the plant occur at the primary cell wall (termed “pit membranes” (Esau, 1965)), that remains intact where xylem conduits of lateral roots join the main axis of nodal roots (McCully & Mallett, 1993; Wang et al., 1994), as well as at the complex plexuses of vascular conduits at the root:shoot junction, and in the nodes of the stem (Shane *et al*., 2000). The pores at these junctions are < 20 nm in diameter and serve to prevent the passage of microbes, soil particles, or embolisms introduced into the system by wounds or pathogen activity (Dong *et al*., 1997).

The impact of perforation plates on the formation, isolation, and repair of embolisms likely has more significant effects on plant water use than their direct resistance to the flow of xylem sap. We anticipated that the short vessel/tall plate phenotype we observed would be less vulnerable to cavitation and contain xylem embolisms more effectively than the long vessel/short plate phenotype. Surprisingly however, genotypes with contrasting perforation plate phenotypes did not display clear differences in vulnerability to cavitation in leaves. Nevertheless, if advancements in methodology could permit more precise measurement of the formation, spread, and repair of embolism formation in other tissues like roots, we may expect to detect differences in these tissues that could not be distinguished in leaves. Specifically, understanding the effects of perforation plates on embolisms in mature root systems would be of interest considering metaxylem of belowground tissue are largest in diameter and length and consequently the most vulnerable to cavitation. The trend of decreasing metaxylem vessel diameter from roots to leaves has been well documented, and the vascular design of most plants typically optimizes water transport in the basal portions of the stem (West *et al*., 1999; Aleman-Sancheschulz *et al*., 2020).

Beyond the direct effects of perforation plates, the observed pleiotropic relationships between perforation plate phenotypes, root elongation rates, root system length and distribution also likely had significant impacts on plant-water relations. The association between the rate of root elongation and the distance between perforation plates along the length of metaxylem vessels suggests that the observed variation in metaxylem structure is driven by differences in cell elongation rates. Differences in root elongation rates consequently appear to contribute to differences in the distribution of root length throughout the soil profile, which has well documented effects on water acquisition.

Specifically, genotypes with the long MXVL phenotype displayed faster rates of root elongation, enabling rapid extension into new soil regions, deeper rooting, and enhanced utilization of deep soil water. This advantage may contribute to enhanced water acquisition, particularly in environments with spatially variable water availability. Phenotypes with deeper rooting also have smaller perforation plates with greater axial conductance, enabling more efficient transport of water originating in deep soil. Conversely, genotypes with short MXVL and slower rates of root elongation display shallower root systems that rely on water acquisition from shallow soil domains. Although we did not observe differences in leaf vessel cavitation in this study, these short MXVL phenotypes also have taller plates which could be linked with reduced root xylem cavitation, which would be advantageous in phenotypes reliant on more temporally dynamic topsoil moisture.

The pleiotropic association of perforation plate phenotypes with root elongation therefore may be an example of synergism between root architecture and root anatomy for water use strategies, as was previously observed in *Phaseolus* taxa, in which greater hydraulic conductance of axial roots is associated with improved drought tolerance in the deep-rooted *P. acutifolius* but is associated with reduced drought tolerance in its more shallow-rooted relative *P. vulgaris* (Strock *et al*., 2021). We propose that the interplay between perforation plate phenotypes and root distribution may represent coordinated strategies to optimize water economies in drought-prone environments. The fact that we observed that the pleiotropic association of xylem vessel length, perforation plate height, and root elongation in the IBM population, which was designed to disrupt genetic linkages arising from gene pool effects, along with the relatively small linkage disequilibrium in maize, an outcrossing species, suggests that such pleiotrophy is genetic rather than the result of evolutionary co-selection. An obvious genetic link can be inferred between the length of xylem vessels and the length of elongating root cells, but the mechanism underlying the association of cell length and perforation plate height is less clear and merits further investigation.

The high heritability of the vessel length/plate height/root elongation pleiotropic syndrome combined with its importance for drought tolerance in maize makes it a potential selection target for the breeding of more drought resilient cultivars. A challenge with many root phenotypes of interest in this context is that they are difficult to phenotype at scale in the field, with some exceptions (e.g. Hanlon *et al*., 2023). Perforation plate height is particularly tedious to phenotype, as plate dimensions are near the resolution of light microscopy. However, vessel length is easier to measure, especially in leaves, where leaf cell length could be rapidly phenotyped in the field with inexpensive pocket USB light microscopes. The low environmental plasticity of this phenotype means that it could be phenotyped in unstressed plants, which facilitates breeding since imposing uniform drought stress in the field is complex and challenging. The relatively simple genetic architecture we report here, with few strong QTL, makes this phenotype amenable to marker assisted selection as well. We propose that this phenotype merits attention as a potential selection target in crop breeding, and that it be investigated in other cereal crops, especially in taxa closely related to maize such as sorghum and millet, but also in wheat, rice, etc.

Initially, we set out to test the hypotheses that (1) simple perforation plates have a significant effect on water transport, (2) intraspecific variation exists for these phenotypes in maize, and (3) this variation affects water use strategies under drought stress. While the first two hypotheses were straightforward to address, the effect of variability in perforation plate structure on water use strategies proved more challenging to disentangle from other pleiotropic effects related to elongation rates. The results of this study shed light on the significance of simple perforation plates in influencing water transport in annual monocots. Observed intraspecific variation in the length of metaxylem vessel elements and the prominence of perforation plates appears to reflect the trade-offs and adaptations to optimize water transport in different ecological niches and could prove useful to breeding programs focused on resistance to drought. The genetic variation for metaxylem structure is directly linked to the efficiency and reliability of water transport within the plant, and their significance should be considered in the context of the specific environmental challenges and opportunities for water use across the distribution of *Zea mays*. Our intensive focus on hydraulics with this work did not afford us the ability to investigate the adaptive significance that root elongation rates have on water capture and use. While this is beyond the scope of the current investigation, it would be worthwhile to more deeply investigate potential synergisms between rates of root elongation and xylem vessel morphology in greater depth.

## Acknowledgments

We thank John Cantolina at the Pennsylvania State University Huck Institutes of the Life Sciences Microscopy Core Facility for assistance with cryo-SEM imaging. This research was supported by the FFAR Crops of the Future and the USDA National Institute of Food and Agriculture and Hatch Appropriations Project #PEN04732. We thank Dr. Shawn M. Kaeppler from U-Wisconsin, Madison, USA for supplying seed for this work.

