## Supplemental Figures for "Xylem perforation plate phenotypes affect water use and drought adaptation in maize (*Zea mays* L.)"

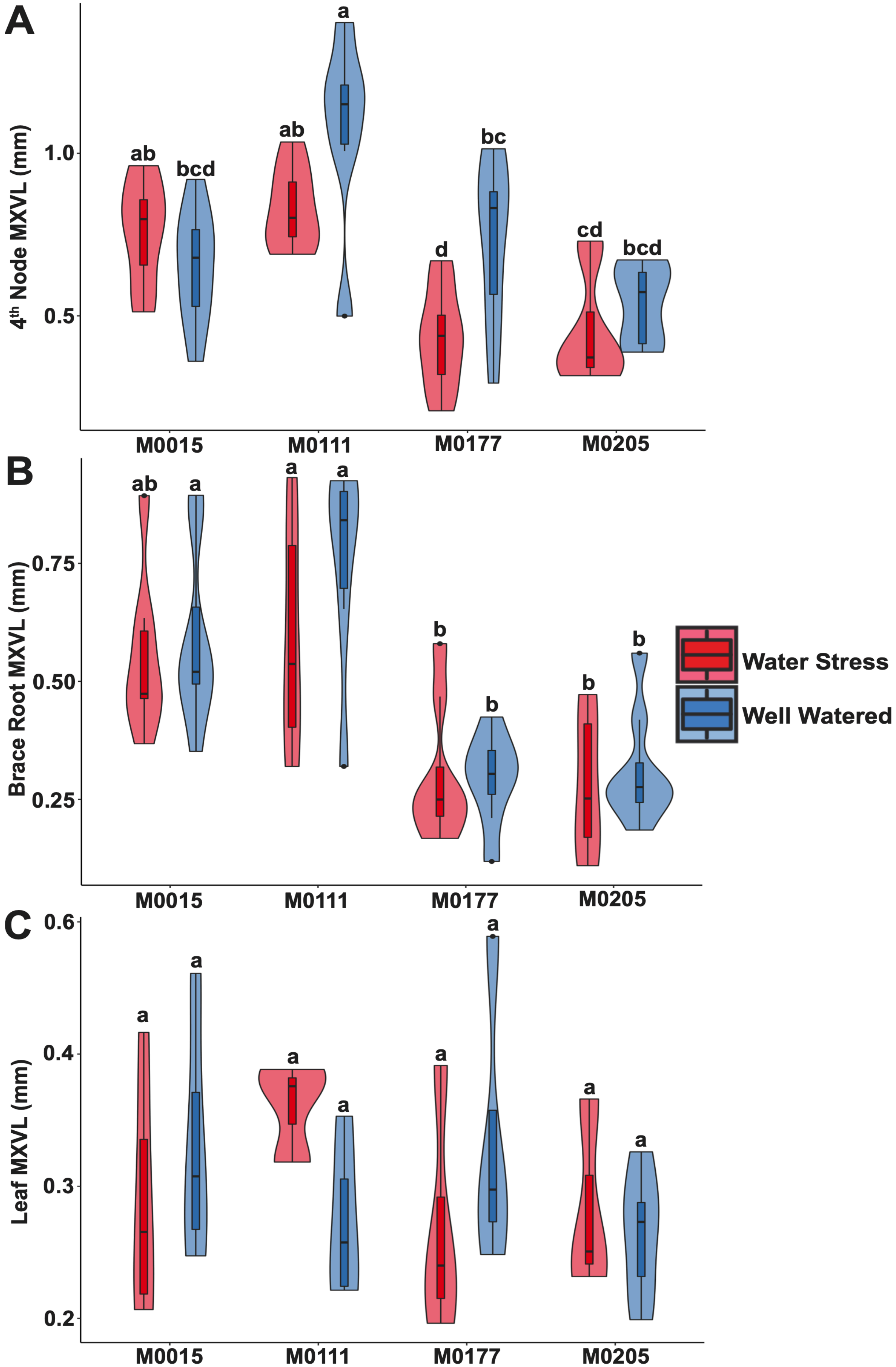


**Supplemental Figure S1.** Violin plots showing the median, interquartile range, 95% confidence intervals, and frequency of MXVL in (A) below-ground 4th node roots, (B) above-ground brace roots, and (C) leaves in the four maize (*Zea mays* L.) genotypes under water stress and well-watered conditions at Graneros, Chile. Letters denote significant phenotypic differences as determined by ANOVA and Tukey HSD (α≤0.05).

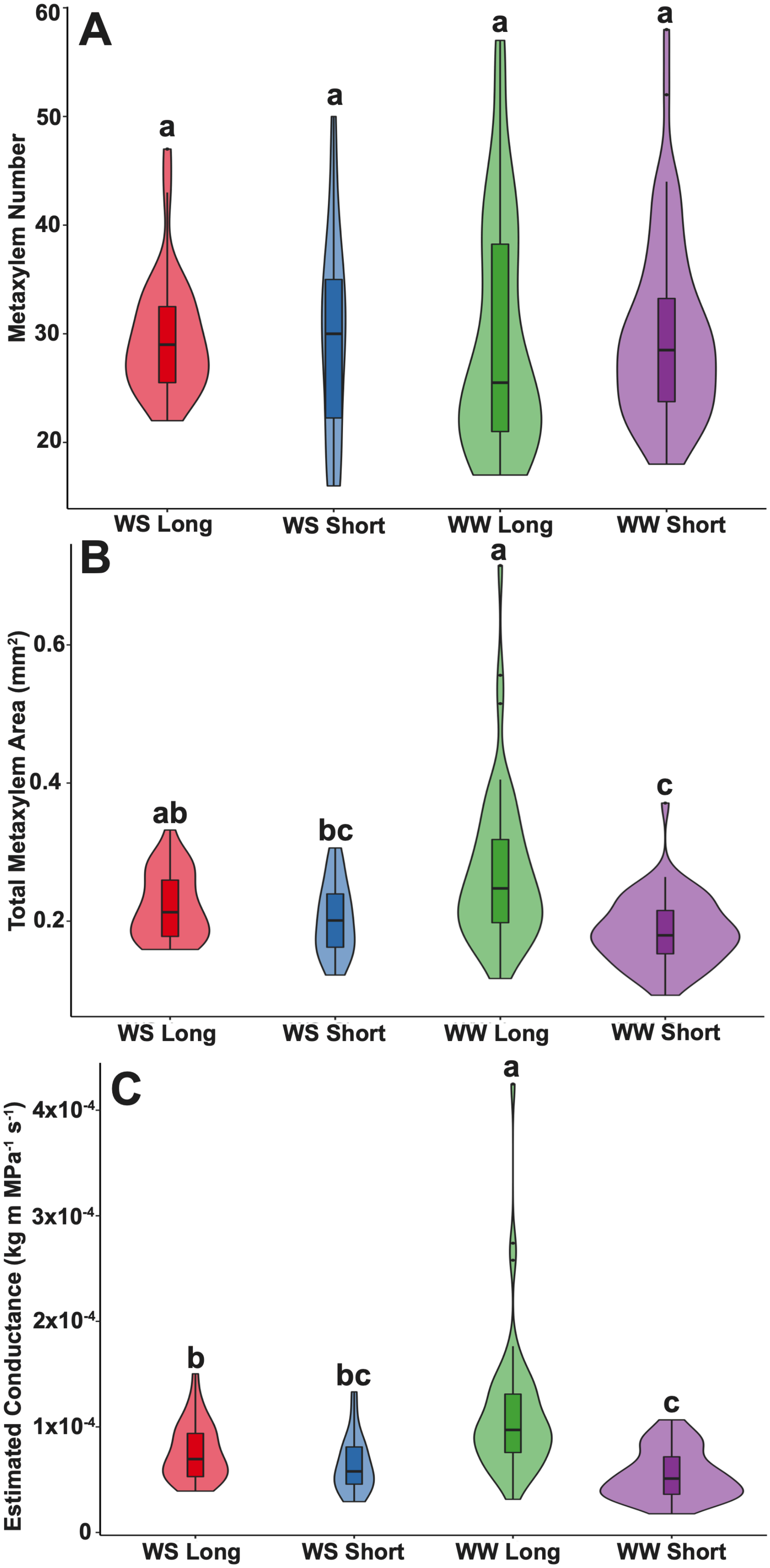


**Supplemental Figure S2.** Violin plots showing the median, interquartile range, 95% confidence intervals, and frequency of metaxylem vessel number (A), total metaxylem vessel area (B), and estimated conductance (C) of below-ground roots in maize (*Zea mays* L.) genotypes with contrasting MXVL under water stress and well-watered conditions in the glasshouse at 42 DAP. Letters denote significant phenotypic differences as determined by ANOVA and Tukey HSD (α≤0.05).

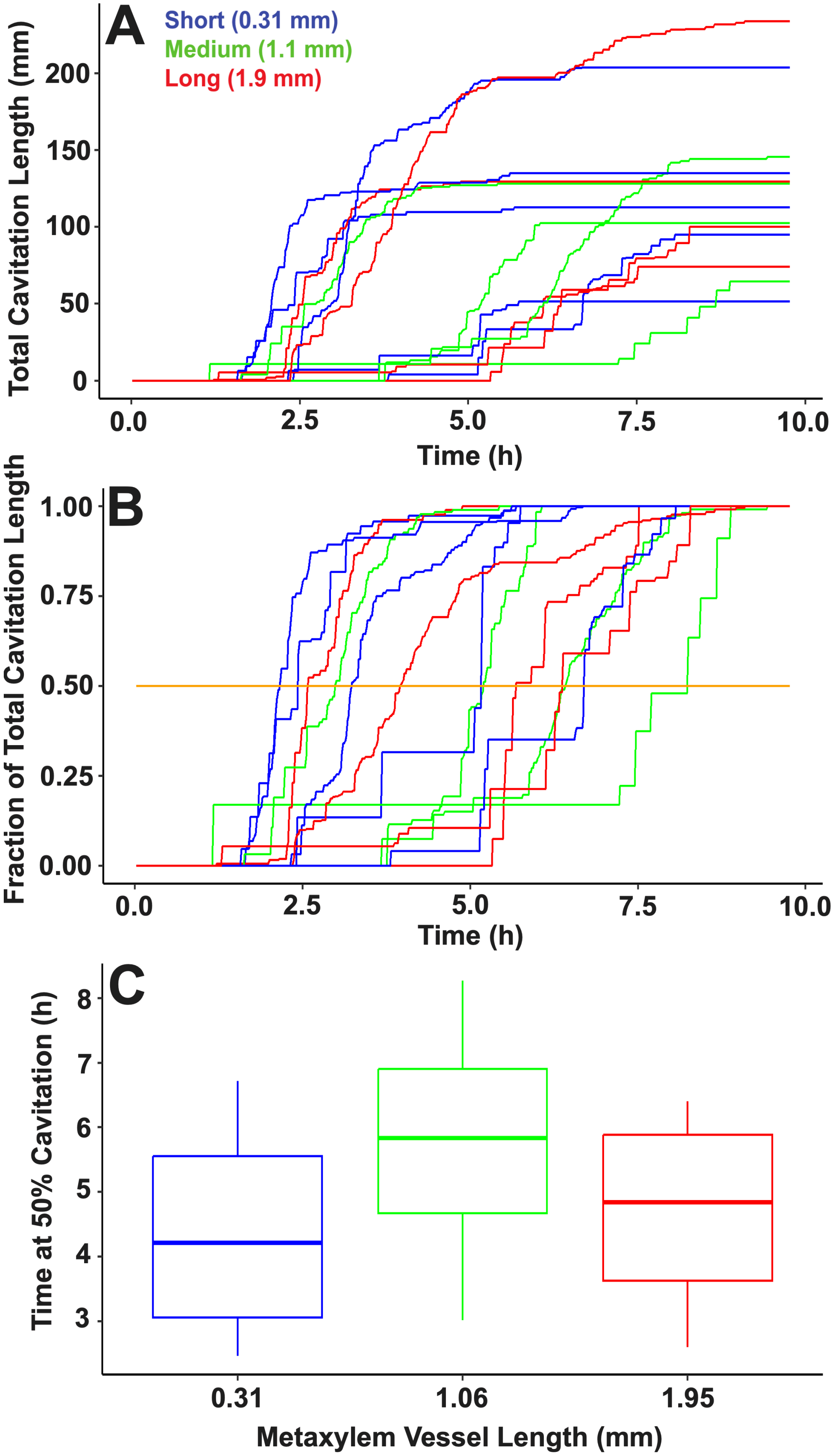


**Supplemental Figure S3.** Metaxylem vessel length has little effect on cavitation time in maize (*Zea mays* L.) leaves. Running total of cavitation event length (mm) in maize leaves with long (red), intermediate (green), and short (blue) metaxylem vessel lengths (A). Each line represents cavitation of a single leaf. Cavitation running totals normalized to 100% and displayed with a threshold value (orange) of 50% (B). Time at 50% total cavitation for various metaxylem vessel lengths (C). No significant difference was observed for time at 50% total cavitation between metaxylem vessel lengths at $p \leq 0.05$ according to the Wilcoxon signed rank test.

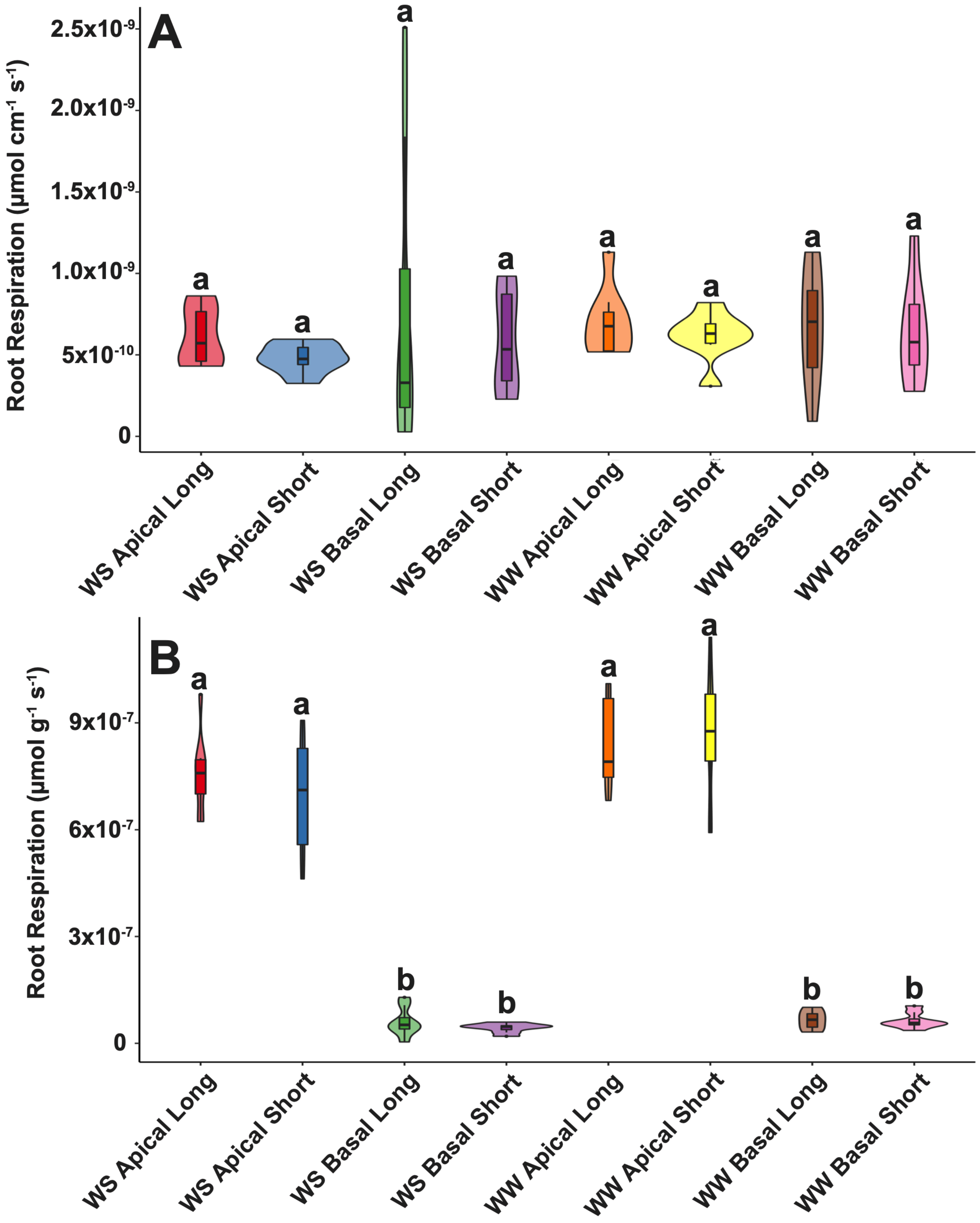


**Supplemental Figure S4.** Violin plots showing the median, interquartile range, 95% confidence intervals, and frequency of root respiration per length of root (A) and root respiration by mass (B) of apical and basal root segments collected from maize (*Zea mays* L.) genotypes with contrasting MXVL under water stress and well-watered conditions in the glasshouse at 31 DAP. Letters denote significant phenotypic differences as determined by ANOVA and Tukey HSD ($\alpha \leq 0.05$).

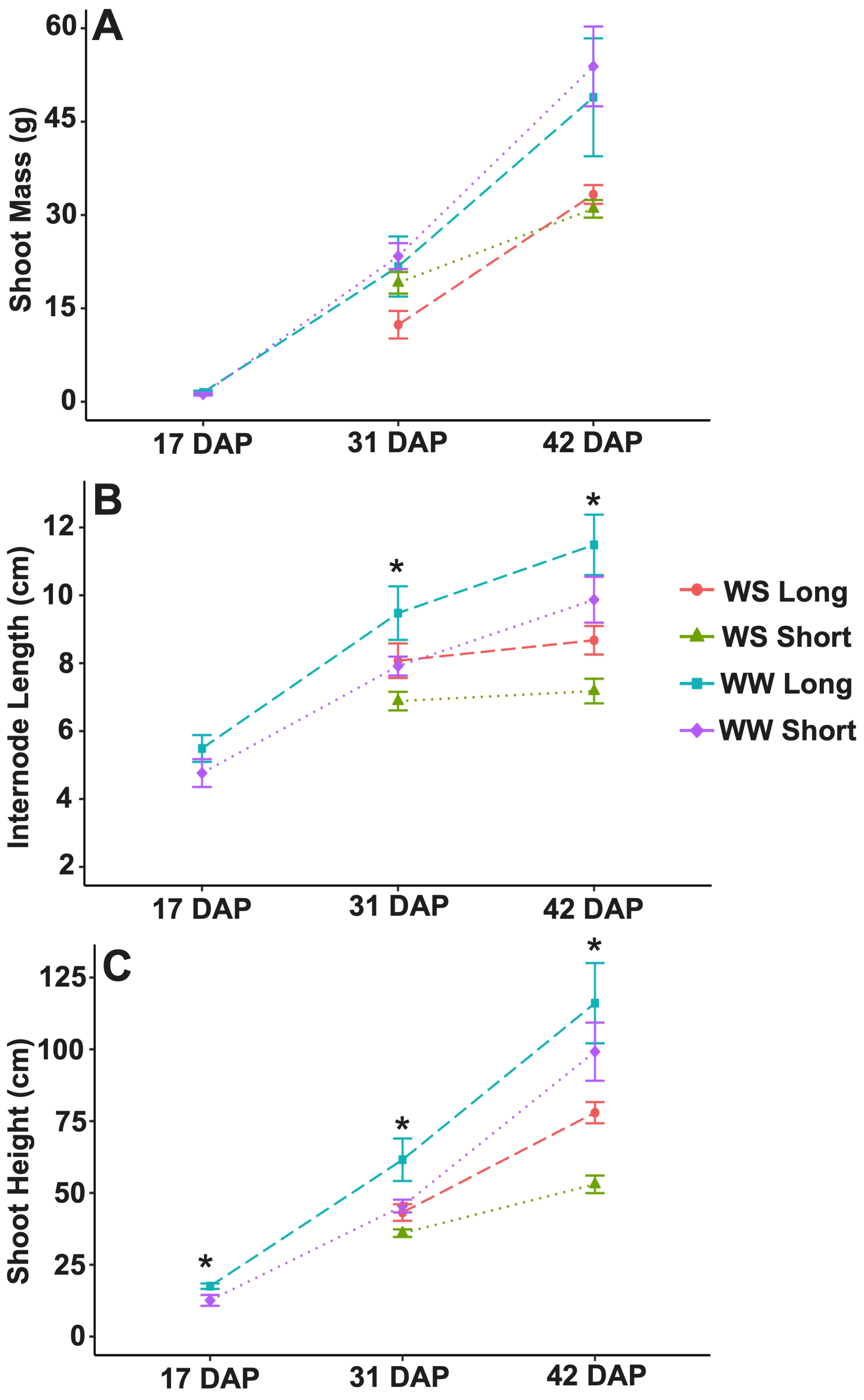


**Supplemental Figure S5.** Dry shoot mass (A), internode length (B), and shoot height (C) of maize (*Zea mays* L.) genotypes with contrasting MXVL under water stress and well-watered conditions in the glasshouse at 17, 31, and 42 DAP. Lines represent mean values (N = 8) and error bars are ± SEM. A significant effect of phenotype at each timepoint as determined by ANOVA ($\alpha \leq 0.05$) is denoted by an asterisk.
